# A bacterial pan-genome makes gene essentiality strain-dependent and evolvable

**DOI:** 10.1101/2022.02.08.479609

**Authors:** Federico Rosconi, Emily Rudmann, Jien Li, Defne Surujon, Jon Anthony, Matthew Frank, Dakota S. Jones, Charles Rock, Jason W. Rosch, Christopher D. Johnston, Tim van Opijnen

## Abstract

Many bacterial species are represented by a pan-genome, whose genetic repertoire far outstrips that of any single bacterial genome. Here we investigate how a bacterial pan-genome might influence gene essentiality, and whether essential genes that are initially critical for the survival of an organism can evolve to become non-essential. By using Tn-Seq, whole-genome sequencing, and RNA-Seq on a set of 36 clinical *Streptococcus pneumoniae* strains representative of >68% of the species’ pan-genome, we identify a species-wide ‘essentialome’ that can be subdivided into universal, strain-specific and accessory essential genes. By employing ‘forced-evolution experiments’ we show that specific genetic changes allow bacteria to bypass essentiality. Moreover, by untangling several genetic mechanisms we show that gene-essentiality can be highly influenced and/or dependent on: 1) the composition of the accessory genome; 2) the accumulation of toxic intermediates; 3) functional redundancy; 4) efficient recycling of critical metabolites; and 5) pathway rewiring. While this functional characterization underscores the evolvability-potential of many essential genes, we also show that genes with differential essentiality remain important antimicrobial drug target candidates, as their inactivation almost always has a severe fitness cost *in vivo*.

## Introduction

No single gene operates in isolation; instead, genetic elements within a genome make up an elaborate network of interacting components that operate in concert to generate a phenotype. While some genes in a genome are essential for growth and survival under any circumstance most genes are non-essential (i.e. dispensable) and their inactivation may have little to no effect on growth^1^. However, their degree of dispensability may depend on the genetic background and/or the composition of the environment. For instance, deletion of some genes will often not trigger a defect if another functionally redundant gene remains. Moreover, genes in a pathway that generate a particular building block, such as Tryptophan, may be dispensable in the presence of the amino acid, but important, or even essential for growth in its absence. In contrast, essential genes are critical for maintaining functionality under any circumstance. This is reflected in their high level of conservation within and often across species, limited genetic diversity, and high and stable expression levels^2–4^. Consequently, essential genes are considered rigid, largely immutable key components to an organism’s survival, which makes them attractive as antimicrobial targets^5, 6^.

New insights are challenging this unyielding essential-gene concept. For instance, large scale experiments in different yeast species have shown that SNPs, loss and gain of DNA sequences, and changes in expression of other genes in the genome can act as mechanisms that suppress gene essentiality^7–10^. Additionally, many species, especially within bacteria, are defined by a pan-genome. This means that different strains may contain distinct sets of genes, resulting in a species with a genetic repertoire that far exceeds that of any single genome. Moreover, a limited number of comparisons using ordered knockout arrays and Tn-Seq^11, 12^ in *E. coli* and *Pseudomonas aeruginosa* experimentally demonstrate that essentiality is sometimes strain-specific^13, 14^. These data indicate that once a gene is essential, it does not necessarily need to remain so. Instead, it suggests that essentiality is a fluid concept likely influenced by both the environment and the genetic background^1, 7^.

The human bacterial pathogen *Streptococcus pneumoniae* is one such species with a large pan-genome. *S. pneumoniae* is a major cause of community-acquired pneumonia, meningitis, and acute otitis media and despite the introduction of several vaccines, remains one of the leading bacterial causes of mortality worldwide^15, 16^. Comparative sequence analysis has shown that, on average, a genome contains ∼2100 genes, while the entire species contains >4000 genes^17, 18^. Moreover, we have shown that two randomly selected strains can differ in genomic content by hundreds of genes, with important phenotypic consequences for antibiotic sensitivity as well as how (antibiotic-mediated) stress is overcome (i.e., which genes and pathways are involved) ^2, 19, 20^. By utilizing Tn-Seq, we have shown that TIGR4, a common lab strain that was originally isolated from a patient with invasive pneumococcal disease^21, 22^, contains >250 essential genes, many of which have been experimentally confirmed. Furthermore, CRISPRi experiments indicate that many essential genes in TIGR4 and D39 overlap^23^. However, a Tn-Seq analysis we performed in three strains (TIGR4, D39, and Taiwan-19F)^2, 19^ suggests that different essential genes exist in *S. pneumoniae*; i.e., essential genes that are non-essential in at least one strain.

In this study, we assemble a detailed functional genomics dataset for 36 *S. pneumoniae* strains covering >68% of the species’ pan-genome. To improve accuracy, most genomes were re-sequenced using single-molecule real-time (SMRT) sequencing (PacBio) to produce long read single contig assemblies, RNA-Seq was performed on all strains, and Tn-Seq on 21 strains within the collection. We show that *S. pneumoniae* contains 206 universally essential genes (those that are present and essential in all strains), 186 strain-specific essential genes (those that are always present but not always essential) and 128 accessory essential genes (those that are essential when present). We use our dataset to show that the different essential-gene types have different characteristics including different expression levels and fitness effects. Furthermore, we use ‘forced-evolution’ experiments combined with whole-genome sequencing to determine how likely it is that a ‘solution’ exists for an essential gene to become non-essential. We find that it is hard for a universal gene to lose its essentiality. In contrast, strain-specific and accessory-essentials can switch between essential and non-essentiality due to changes in the genetic background, including specific gene deletions, bypass mutations, and gene-expression changes. Importantly, we show that while essentiality may be dependent on the genetic background, those genes demonstrating essentiality only in certain contexts remain important antimicrobial drug target candidates, as their deletion almost always has a fitness cost *in vitro* and *in vivo*.

## Results

### A representative strain collection of the S. pneumoniae pan-genome

A Pan-Genome (PG) strain collection was assembled consisting of 36 *S. pneumoniae* strains representing 16 different capsule serotypes (Supplementary Table 1). Six are commonly used experimental strains including D39 and TIGR4^21, 24^, while 30 strains are part of a larger (>350) invasive *S. pneumoniae* surveillance study strain collection, isolated at a hospital in Nijmegen, the Netherlands^25^. Notably, these 30 strains consist of 15 genetically similar pairs, which should display similar phenotypes among pairings. Of the 36 total strains, 33 were re-sequenced with PacBio to obtain a single contig for each strain (Bioproject PRJNA514780).

To bioinformatically test whether the PG-collection reflects the *S. pneumoniae* pan-genome, we created a ‘pan-genome study group’ consisting of our 36-strain PG-collection, 23 reference genomes ^26^, 111 strains from several worldwide surveillances programs (30 from Massachusetts, US, 30 from Maela, Thailand, 30 from Southampton, UK, and 21 from Malawi), and 38 isolates obtained before 1974 (Supplementary Table 2)^27–30^. Three different pan-genome analysis tools^31–33^ were used to estimate the average number of genes/strain at ∼2100 genes, the size of the pan-genome at >4100 genes (all genes in the species), while the core-genome consists of at least 1349 genes (present in all strains), which is similar to previous estimates^17, 18^ (Fig. 1A) (Supplementary Table 2 and 3). Our 36-strain PG-collection covers >68% of genetic content across the pan-genome, and scatters evenly through a maximum-likelihood phylogenetic tree (Fig. 1B), indicating that the PG-collection is indeed representative of the *S. pneumoniae* pan-genome.

**Figure 1:**
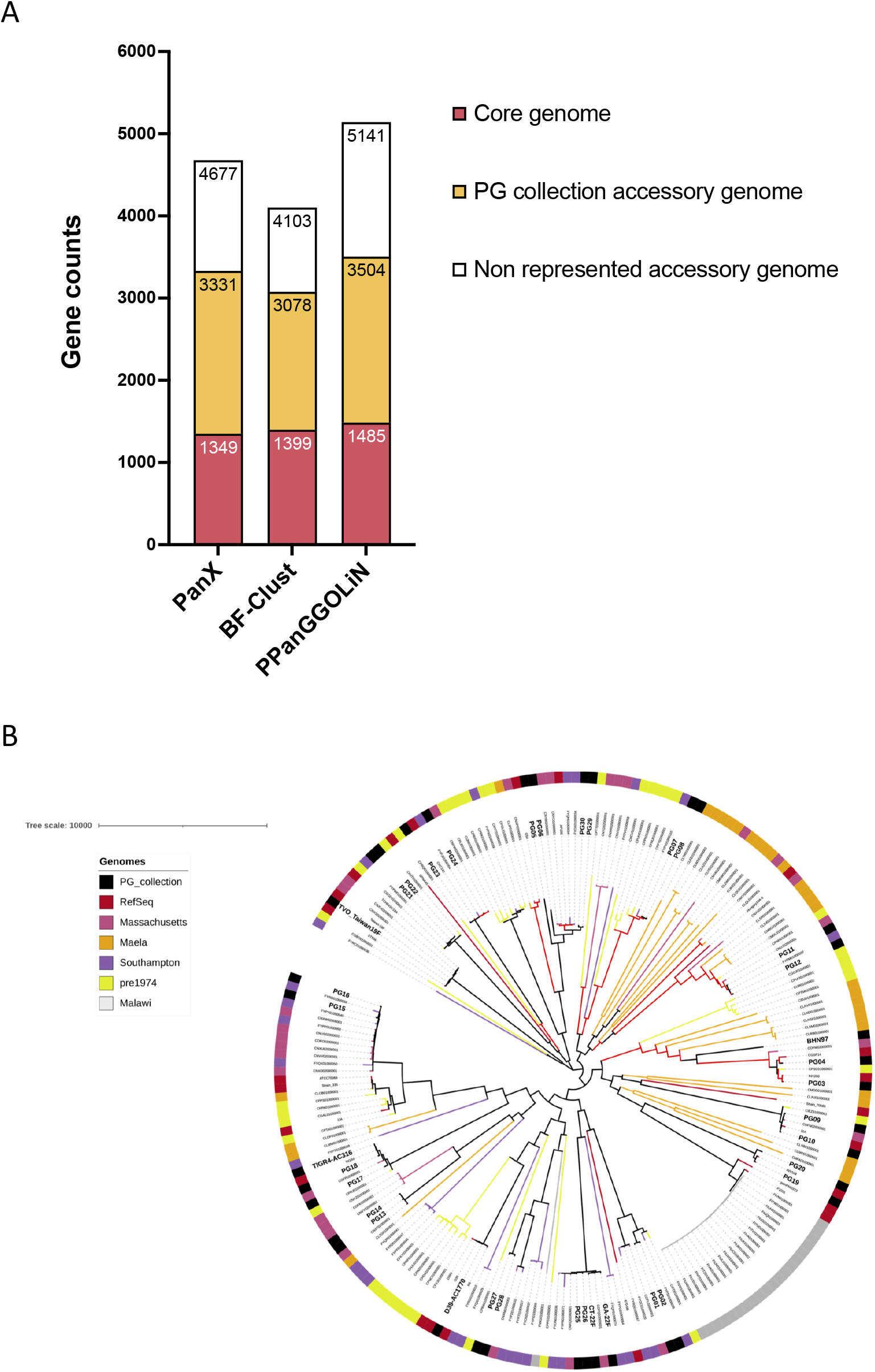
*Streptococcus pneumoniae* <u>P</u>an-<u>G</u>enome collection genetic content and phylogenetic diversity. A) Number of gene clusters belonging to the core and accessory genome for the 208 strains study group determined by three different methods (PanX, BF-Clust, PPanGGOLiN). Orange represents how much of the accessory genome is present in the 36 strains of the PG collection. B) Maximum-likelihood phylogenetic tree based on core genes SNPs of the 208 strains in the pan-genome study group. Labels from the strains selected for our PG collection are in bold and have a larger font size. Black outer strip color symbol and black tree branches also indicate the PG collection strains.

### Universal and strain-dependent gene essentiality

To determine if gene-essentiality is dependent on a strain’s genetic background we employed Tn-Seq analysis. Transposon library construction in *S. pneumoniae* relies on the bacterium’s natural competence/transformability, the efficiency of which is affected by different genetic factors, such as capsule serotype^34–36^. Transformation efficiency was variable (Supplementary Table 1), leading to the successful construction of transposon libraries in 21 out of 36 strains (Fig. 2A). Tn-Seq data analysis can predict gene essentiality based on library saturation (i.e., percentage of occupied TA sites by the *mariner* transposon) and the lack of insertions in a gene. Out of 21 libraries, 17 had a saturation >35% enabling high-confidence essentiality predictions (Fig. 2A). The binomial method from the TRANSIT package was used to make essential-gene calls^37, 38^, which categorizes genes as ‘Essential’, ‘Uncertain’, or ‘Non-Essential’. For strains with >35% saturation, the number of essential genes range from 274 (PG29) to 379 (PG02) (Supplementary Table 4). Compared to previous findings there was ∼90% overlap for essential gene calls made for common lab strains TIGR4, D39, and R6^23, 39–42^. From the high confidence ‘essential’ category, we define the ‘essentialome’ of *S. pneumoniae* as consisting of 520 genes that are essential in at least one strain. Using gene cluster comparisons of binomial predictions across strains, we further subdivide the essentialome into: **1)** 206 <u>Universal essentials</u>: core genome genes that are present and essential in each strain; **2)** 186 <u>Strain-dependent essentials</u>: core genome genes that are present in all strains but only essential in some; and **3)** 128 <u>Accessory essentials</u>: accessory genome genes predicted as essential in the strains where they are present (Fig. 2B, C and Supplementary Table 5). This categorization of strain-dependent and accessory essentials suggests that for a substantial number of genes in *S. pneumoniae* essentiality is genetic background-dependent.

**Figure 2:**
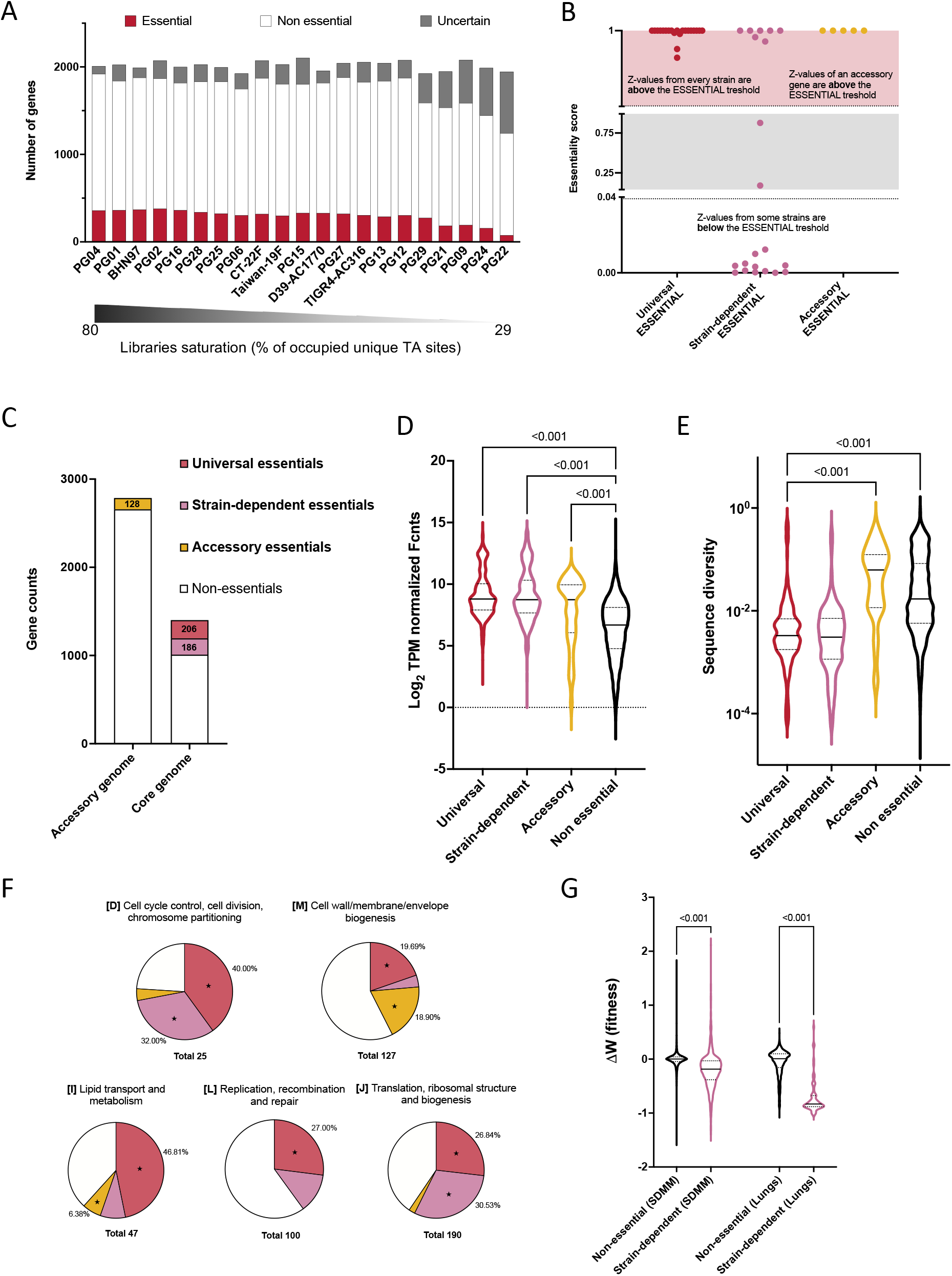
Essential genes of *Streptococcus pneumoniae* PG collection. A) Number of genes called Essential, Non-Essential, or Uncertain by TRANSIT Binomial method in the 21 strains with mutant libraries. The bar below shows the Tn-Seq libraries saturation measured as the percentage of the total unique TA sites in the genome occupied by the *mariner* transposon. B) Figure represents 3 representative genes showing the criteria used to classify genes as “Universal essentials”, “Strain-dependent essentials”, and “Accessory essentials”, based on the Binomial method. Each dot represents the essentiality score obtained for a gene in a specific strain. An essentiality score above 0.9916 categorizes a gene as essential and below 0.0424 as non-essential, as determined by a FDR 5% approach. C) Number of genes assigned to the different classes. D) Gene transcript abundances obtained by RNAseq collected from early exponential phase cultures of the 36 PG collection strains growing in THY medium. Graph shows genes grouped by the essentialome classification depicted in B. Black lines inside each violin plot represent the median, and dotted lines the first and third quartiles. Tukey’s corrected one-way ANOVA comparison *p*-values between non-essential and the other three categories are shown. E) Sequence diversity of *S. pneumoniae* genes split into the different essentialome classes and measured using the diameters of the clusters obtained by BF-Clust. Black lines inside each violin plot represent the median, and dotted lines the first and third quartiles. The *p*-values shown resulted from a Kruskal-Wallis test comparison. Strain-dependent versus Accessory and Non-essentials classes comparison presented the same *p*-value (<0.001). F) Functional categories enriched in Universal, Strain dependent, and Accessory essentials. Each chart represents a COG category and the fraction of genes belonging to each essentialome class. Color code is the same as C, D, and E panels. Asterisks indicate enrichments of essentialome class within COG category with an adjusted *p*-value< 0.05. G) Distribution of gene fitness differences calculated by Tn-Seq for each strain, using transposon mutants’ libraries growing in medium SDMM (left plots) and for strain TIGR4 libraries recovered from mouse lungs (right plots)^42^. Graph shows genes grouped by essentialome class. Tn-Seq calculates fitness only for non-essential genes; thus, the “Strain-dependent” distribution plot shows the fitness differences observed for only those strains where the genes are non-essential. Black lines inside each violin plot represent the median, and dotted lines the first and third quartiles. The *p*-value obtained from Tukey’s corrected one-way ANOVA comparison between the two distributions is shown.

### Universal and strain-dependent essential gene characteristics

Essential genes may have characteristics that distinguish them from non-essential genes. For instance, an essential gene’s expression is often higher, and much less affected by environmental changes than the average non-essential gene^1, 2, 4, 43^. We assessed whether the identified essentialome genes have different characteristics compared to non-essential genes: **1)** RNA-Seq was performed on the 36-strain PG collection grown in rich medium, which shows that the universal, strain-dependent and accessory essential genes tend to be more highly expressed compared to non-essential genes (Fig. 2D); **2)** Essential genes tend to be more conserved and contain less genetic variation^1, 3^. We calculated the sequence diversity for each gene across all 36 strains, which shows that universal and strain-dependent essential genes are less genetically diverse than accessory and non-essential genes (Fig. 2E). Moreover, the higher diversity in accessory essentials may be because many are involved in ‘addiction-like’ systems. Such systems often consist of two or more genes in which one of the genes is only required if the other/s are present, which includes phage repressors, antitoxins from toxin-antitoxin systems, and methylases from restriction-modification systems (Supplementary Table 5). These genes may thus tolerate more mutations than a core genome gene involved in central processes, such as DNA replication; **3)** Functional enrichment analysis shows that universal essential genes are enriched in 5 COG categories (Fig. 2F), all of which support central cellular processes and have been found as enriched in essential gene sets from different species^4, 44, 45^, while strain-dependent and accessory essential genes are enriched for only two of these categories (Fig. 2F); **4)** Since gene essentiality for some genes depends on the strain background, we reasoned that they would likely still trigger a growth defect in the strains where they are not essential. We used Tn-Seq to determine the fitness effect caused by inactivation of these strain-dependent essential genes in rich medium (Supplementary Table 6). Results show that mutants with insertions in strain-dependent essentials, on average, have a significant fitness defect compared to mutants with insertions in non-essential genes (Fig. 2G). Moreover, Tn-Seq data obtained from a mouse lung infection model ^42^ shows that strain-dependent essentials also have a significant defect *in vivo* compared to non-essentials (Fig. 2G). These data highlight that essential genes have characteristics that differentiate them from non-essential genes, and in some cases distinguish the universal, strain-dependent and accessory groups from each other. Importantly, the fitness defect caused by inactivation of strain-dependent essential genes shows that genetic-background mechanisms responsible of essentiality bypass may not be able to entirely compensate for gene loss.

### Strain-dependent essentiality is a fluid and evolvable state

Genes do not operate in isolation; i.e., they interact with other genetic elements and processes within a complex genomic network to generate a phenotype. The existence of genes with variable essentiality underscores that is not an entirely fixed concept; instead genome composition and genetic interactions may make the essentiality concept more fluid. This suggests that at least some essential genes are evolvable and can switch between an essential and non-essential state depending on their network connectivity and/or genetic background. To determine whether different essential genes have different levels of evolvability, we targeted five universal and six strain-dependent essentials in four different strain backgrounds to obtain knockout mutants (Fig. 3A and B). Transformation with a PCR product containing a drug marker fused to ∼2000bp of DNA flanking a non-essential gene of interest will normally, with increasing amounts of DNA, generate an increasing number of transformants in which the targeted gene is knocked out^46, 47^. However, during the entire transformation process additional lesions including SNPs or small deletions may occur elsewhere in the genome. We reasoned that by using more DNA we could increase the chance of recovering an essential gene-knockout due to simultaneously occurring mutations that bypass and alleviate the loss of the essential gene (Fig. 3A, B, and C).

**Figure 3:**
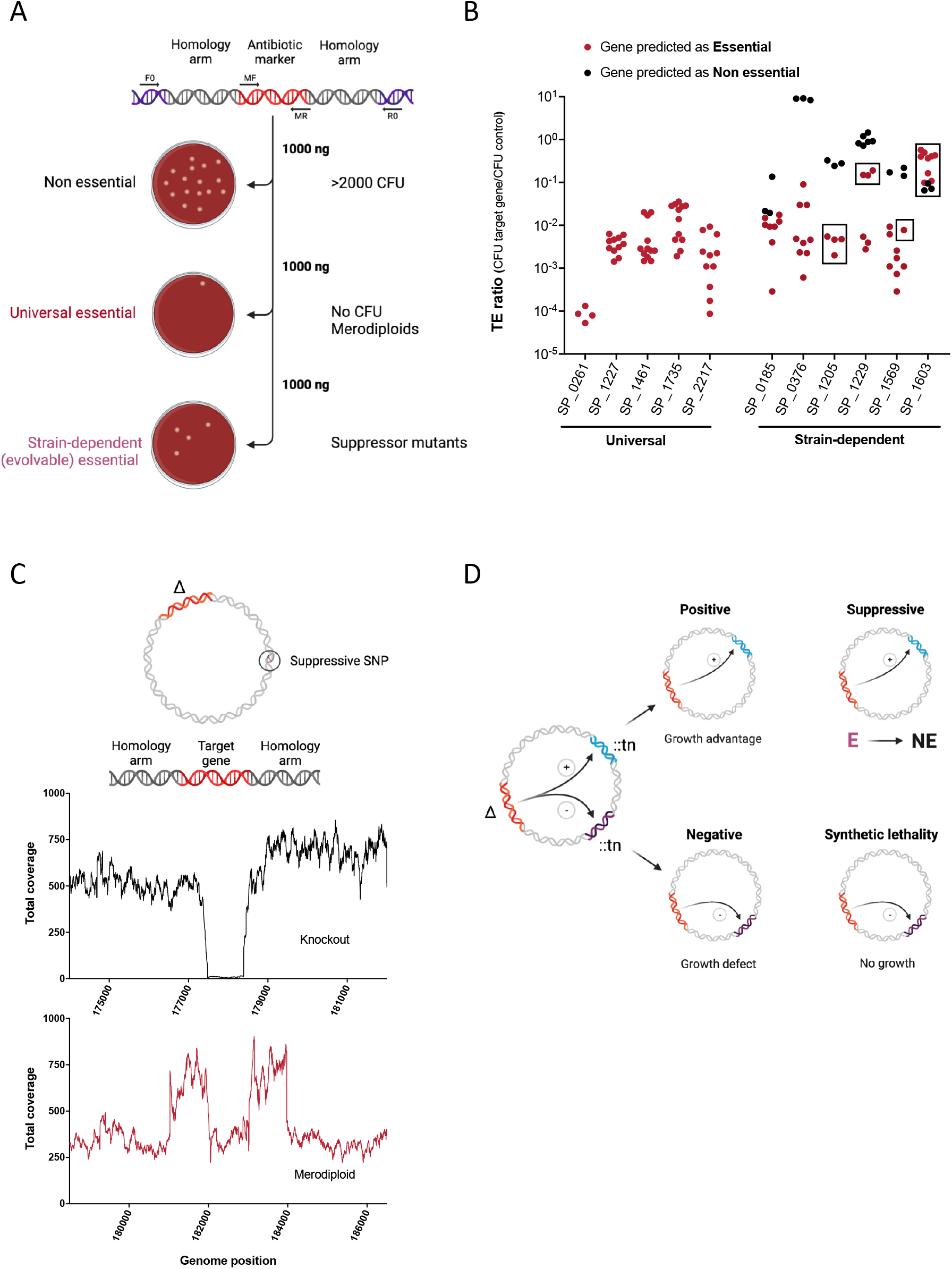
Methodology for essentiality validation, evolvability, and genetic interactions identification of essential genes. A) Strategy deployed to obtain knockouts in different genes and the expected results for each essentialome class. The amount of PCR knockout product used (1000 ng) was 20 times higher than routinely used. B) Validation of gene essentiality. Transformation efficiency (TE) ratio is the division of CFU obtained after transforming 1000 ng of a PCR product targeting an essential gene divided by the CFU obtained after transforming 1000 ng of a PCR product that inserts in a neutral genome region. The graph shows the TE ratios after targeting five universal and six strain-dependent essential genes in four different strain backgrounds by triplicate. Each dot represents one of these replicates, black dots indicate strains where the targeted gene is non-essential, and red dots where the gene is essential. Squares indicate the transformation experiments with successful knockout recoveries. C) Knockout confirmation and suppressive mutations identification. Whole-genome sequencing analysis (*breseq*) of clones selected from transformations revealed if the clones were true knockouts (targeted gene coverage is near zero, middle panel) or merodiploids (target gene coverage same as average, homology arms coverage twice the average, bottom panel). *Breseq* also identified SNPs in genes putatively involved in gene essentiality suppressive mechanisms. D) Genetic interactions between strain-dependent and non-essential genes. Tn-Seq experiments using libraries constructed in different knockouts identified four different types of interactions: 1) a positive interaction occurs when a double mutant (knockout, “Δ”, plus transposon, “::tn”) presents a growth advantage compared to individual mutants; 2) suppressive interaction occurs when one essential gene becomes non-essential in the context of the knockout; 3) in a negative interaction the double mutant presents a growth defect, and 4) in a synthetic lethality the double mutant is non-viable. Tn-Seq fitness calculation identifies positive and negative interactions while TRANSIT binomial method identifies suppressive and synthetic lethality.

When targeting the five universal essential genes in 4 different strains, the transformation efficiency was generally much lower than when targeting a non-essential gene. However, a surprisingly large number of colonies grew for 18 out of the 20 gene/strain combinations (Fig. 3B). Whole genome sequencing (WGS) was performed on 25 randomly picked colonies and 6 entire plate-populations (Fig. 3C and Supplementary Table 7). This approach revealed that these colonies were, in fact, merodiploids, i.e., bacteria containing genomes in which a wild-type copy of the essential gene is retained^48^. Merodiploidy occurs typically at a low frequency in *S. pneumoniae*, and thus the large amounts of DNA used in the transformation likely accounts for the high merodiploid frequency. Importantly, besides the merodiploids, no other ‘solutions’ were found for these five universal essentials, confirming their essentiality could not be easily overcome.

For the six strain-dependent essential genes, transforming the strain-backgrounds in which based on Tn-Seq data these genes are non-essential, yielded as expected, many transformants (Fig. 3B), and these knockouts were confirmed with WGS (Fig. 3C). Transformation of the strains in which the genes are predicted to be essential showed three types of results: **1)** For genes SP_0185 and SP_0376 a low number of colonies were recovered, which were all merodiploids. This suggests that essentiality for these genes is not easily evolvable or bypassed; **2)** For gene SP_1603/*cmk* a high number of colonies were recovered. WGS confirmed clean knockouts, and showed no additional genetic changes in the genomes, indicating that SP_1603 is, in fact, a non-essential gene; **3)** For genes SP_1229 and SP_1205 a low number of colonies were recovered, consisting of a mix of true knockouts and merodiploids. For instance, in strain PG01 only merodiploids were recovered for gene SP_1229, while in strain PG16, WGS identified the knockout and additional genetic changes that could be circumventing essentiality. In a similar fashion, potentially successful bypass mutations in two different backgrounds were identified for strain-dependent essential SP_1205. These data suggest that strain-dependent essential genes can indeed lose their essentiality through changes in a strain’s background, which are explored in more detail below.

### cmk is a non-essential gene whose inactivation affects growth in a strain-dependent manner

While our forced evolution experiments suggest that SP_1603/*cmk* is a non-essential gene, the role of this gene is not entirely clear, which prompted us to explore why *cmk* was predicted as a potential strain-dependent essential gene. *cmk* has been indicated as an essential gene in *S. pneumoniae* for almost two decades, even though small colonies have been observed after transformation^49^. Additionally, our Tn-Seq data predicts *cmk* to be essential in 12 out of 21 strains (Supplementary Table 4). However, we were able to obtain many transformants in three strain backgrounds where *cmk* was predicted as essential (Fig. 3B); i.e., WGS of 13 clones confirmed 12 true knockouts and only one merodiploid (Supplementary Table 7).

The primary role suggested for *cmk* is the recycling of CMP back to CDP (Fig. 4A). In *E. coli* Δ*cmk* presents a growth defect, accumulates 30-fold more CMP than the wild type, and dCTP pools decrease by half^50^. Here we show that growth of *S. pneumoniae* Δ*cmk* in two of three different backgrounds reveals an extended lag phase (Fig. 4B), which makes the mutant unfit, and may explain why it is lost during transposon library construction and indicated as essential. Growth curves also highlight strain-dependent differences, with strain PG04 having the most severe defects (Fig. 4B). These phenotypes could be caused by a decreased replication rate provoked by a dCTP shortage. Then, adding excess uracil and/or cytidine to the growth medium could potentially restore growth, however this had no effect on growth in *S. pneumoniae* (Extended Data Fig. 1A). Alternatively, CMP accumulation may be negatively affecting pathways or reactions where CMP is a product. In *S. pneumoniae*, genes SP_1273 and SP_1274 generate CMP from CDP-choline and are involved in decorating teichoic acid (TAs) with choline^51, 52^. While CMP accumulation could thus affect TA choline decoration, the addition of 10x more choline to the medium also did not compensate growth (Extended Data Fig. 1A).

**Figure 4:**
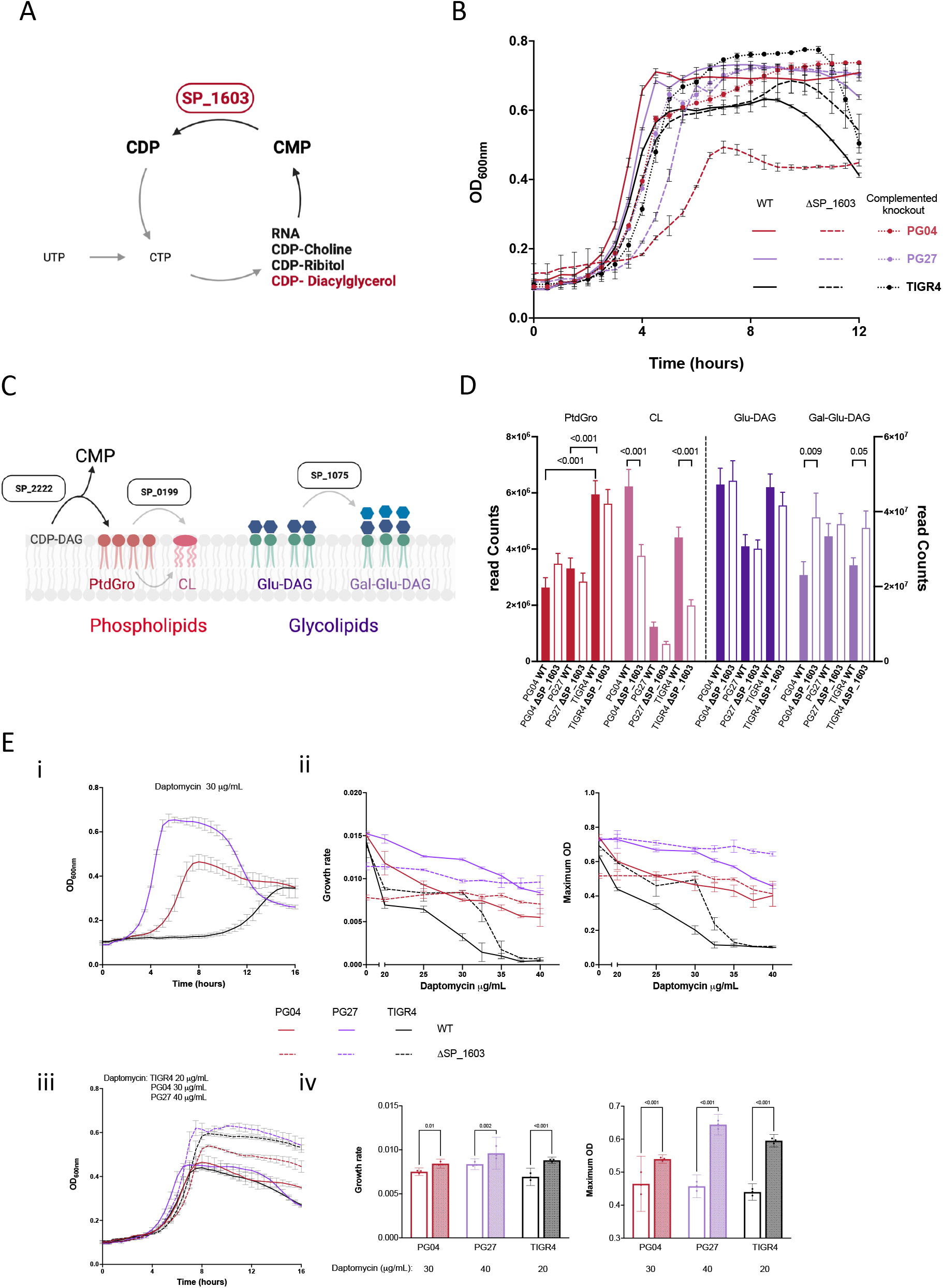
*cmk* deletion alters polar membrane lipid composition. A) Schematic representation of SP_1603 function in CMP recycling. B) Growth phenotype of wild type and derived SP_1603 knockouts in medium SDMM. C) Genes involved in the synthesis of the phospholipids phosphatidylglycerol (PtdGro) and cardiolipin (CL) and the glycolipids glucosyl-diacylglycerol (Glu-DAG) and galactosyl-glucosyl-diacylglycerol (Gal-Glu-DAG). D) Membrane polar lipid composition of different strains and their SP_1603 derived knockouts. Bar graph represents the average read Counts ± SD (n=2) of the different C-14 labeled polar lipids in the membrane of the wild type strains PG04, PG27, and TIGR4 (filled bars) and derived SP_1603 knockouts (open bars). Red bars represent PGP amounts, pink CL, dark violet Glu-DAG and light violet Gal-Glu-DAG. *P*-values depicted are from an Ordinary One-Way ANOVA comparison between wild type and knockout lipid levels. E) Growth phenotype of the different strains grown in the presence of the drug daptomycin. i) Wild types growth curves in the presence of 30 μg/mL of daptomycin. ii) Growth rate and maximum OD of the different strains growing in increasing concentrations of daptomycin. iii) Growth curves of the different strains at 20 (TIGR4), 30 (PG04), and 40 (PG27) μg/mL of daptomycin. iv) Growth rate and maximum OD of the curves depicted in iii. *P*-values depicted are from an Ordinary One-Way ANOVA comparison between wild type and knockout growth in the same condition. Data shown are the results from a single experiment with three replicates per condition (three different wells, same 96-well plate). Independent repetition of the experiment (>3, different plates, different days) showed similar results.

CMP is also a product of the reaction catalyzed by PgsA (SP_2222), an enzyme required for phosphatidylglycerol (PtdGro) synthesis (Fig. 4C). Mutations in *pgsA* cause lower levels of PtdGro and cardiolipin (CL) at the membrane and confer daptomycin resistance in four distinct Gram-positive species^53, 54^. Additionally, in *S. pneumoniae* ΔSP_0199 does not produce CL and is more sensitive to daptomycin^19^. CMP accumulation, mediated by Δ*cmk*, may thus cause membrane lipid composition changes, and changes in the susceptibility to daptomycin. To explore this, we first examined the membrane polar lipid composition by acetate-labelling and TLC, and second the growth in the presence of daptomycin of the different strains (Fig. 4D and E, Extended Data Fig. 1B).

These data show that polar lipid composition is strain specific. For instance, TIGR4 PGP levels are almost twice as high as the phospholipid levels of the other two strains. Due to the role of PtdGro in daptomycin sensitivity, this difference may explain why TIGR4 is the most sensitive to daptomycin of the three wild type strains (Fig. 4E)^55^. All knockouts present a decrease in CL levels and increase in Gal-Glu-DAG levels, although changes in PG27 are modest. For the wild type strains increasing concentrations of daptomycin cause an extended lag-phase, diminution of the growth rate, and lower maximum ODs (Fig. 4E). In contrast, daptomycin only mildly affects the growth phenotype of the knockouts. Importantly, previous Tn-Seq data from TIGR4 shows that ΔSP_1075, the gene responsible for Gal-Glu-DAG synthesis (Fig. 4C), is highly sensitive to daptomycin^19^. The decreased sensitivity of Δ*cmk* may thus be at least partially explained by the increase in Gal-Glu-DAG levels. These data highlight how differences in lipid composition exist between strains and how there is thus not one perfect way of building a lipid membrane. How cmk deletion affects Gal-Glu-DAG levels is unclear since there is no direct metabolic link between CMP and glycolipid synthesis. However, since UTP is the precursor to CTP Δ*cmk* could result in lower UTP pools. Then, this pathway would need to be ramped up in the absence of recycling via *cmk* product, and this could explain the impact on glycolipid synthesis. Importantly, results show how something as seemingly simple as CMP accumulation can affect lipid composition. And thus, while *cmk* is important for growth, but not essential, it illustrates how its effects on the processes it is connected to can vary significantly depending on the genetic background.

### S. pneumoniae accessory essentials interact with core genome genes

The category of accessory essentials includes many genes part of genetic addiction modules^56^, which, as explained above, includes gene-pair modules where the product of one of the genes protects cells against the toxicity of the other. For instance, the antitoxin coded by SP_1809 is essential because it is needed to counteract the toxic effect of gene SP_1810. In the proposed mechanism of this toxin-antitoxin pair, SP_1809 requires the Clp protease subunit ClpX (SP_1569) to degrade the toxin produced by SP_1810^57^. Indeed, *clpX*/SP_1569 is essential only in the PG collection strains which harbor the SP_1809-1810 gene pair (Supplementary Table 4). Moreover, we show that the genetic loss of the toxin/antitoxin system allows for the recovery of a *clpX*/SP_1569 knockout from PG27, a strain where *clpX* is essential (Fig. 3B and Supplementary Table 7). This interaction between *clpX*, a core genome gene, and this toxin/antitoxin system represents an example of how the accessory genome can influence essentiality of core genes.

The most enriched functional category of accessory essentials are genes involved in cell wall, membrane and envelope biogenesis (Fig. 2F), which can be explained by the high diversity of essential capsule type-specific biosynthesis genes. The capsule is one of *S. pneumoniae’s* most important cellular structures and is essential for immune escape and *in vivo* survival^58^. This selective pressure may explain the high genetic diversity in capsules, which currently stands at 100 different serotypes^59^. *S. pneumoniae’s* capsule synthesis genes are mostly organized in a large operon 20 - 30 kilobases long^60^. While most of these genes are accessory essentials (Fig. 5A, Supplementary Table 5), the sugar transferases, or phosphoglycosyl transferases^61^ responsible for attaching the first capsule residue to the lipid carrier undecaprenyl phosphate (UP), are non-essential in every strain (Supplementary Table 4). In D39 missense and nonsense point mutations in the sugar transferase can relieve essentiality of downstream capsule genes^62, 63^. To determine whether this applies across the pan-genome, we constructed knockout mutants of the sugar transferase in four different strains (D39, PG06, Taiwan19F, and TIGR4) and subsequently introduced transposon libraries in the background of this query gene. Tn-Seq data shows that within the sugar transferase query gene background, the essentiality of the downstream capsule genes is indeed suppressed (Fig. 5B). As previously proposed, the most plausible explanation for this suppression is that the essential molecule UP gets trapped if the downstream capsule genes are non-functional, subsequently affecting its recycling and the synthesis of peptidoglycan and teichoic acids. However, by blocking the attachment of the initial UDP-sugar residue to UP (e.g. by knocking out the sugar transferase, Fig. 5), the lethal dead-end lipid carrier trapping can be prevented^60^.

**Figure 5:**
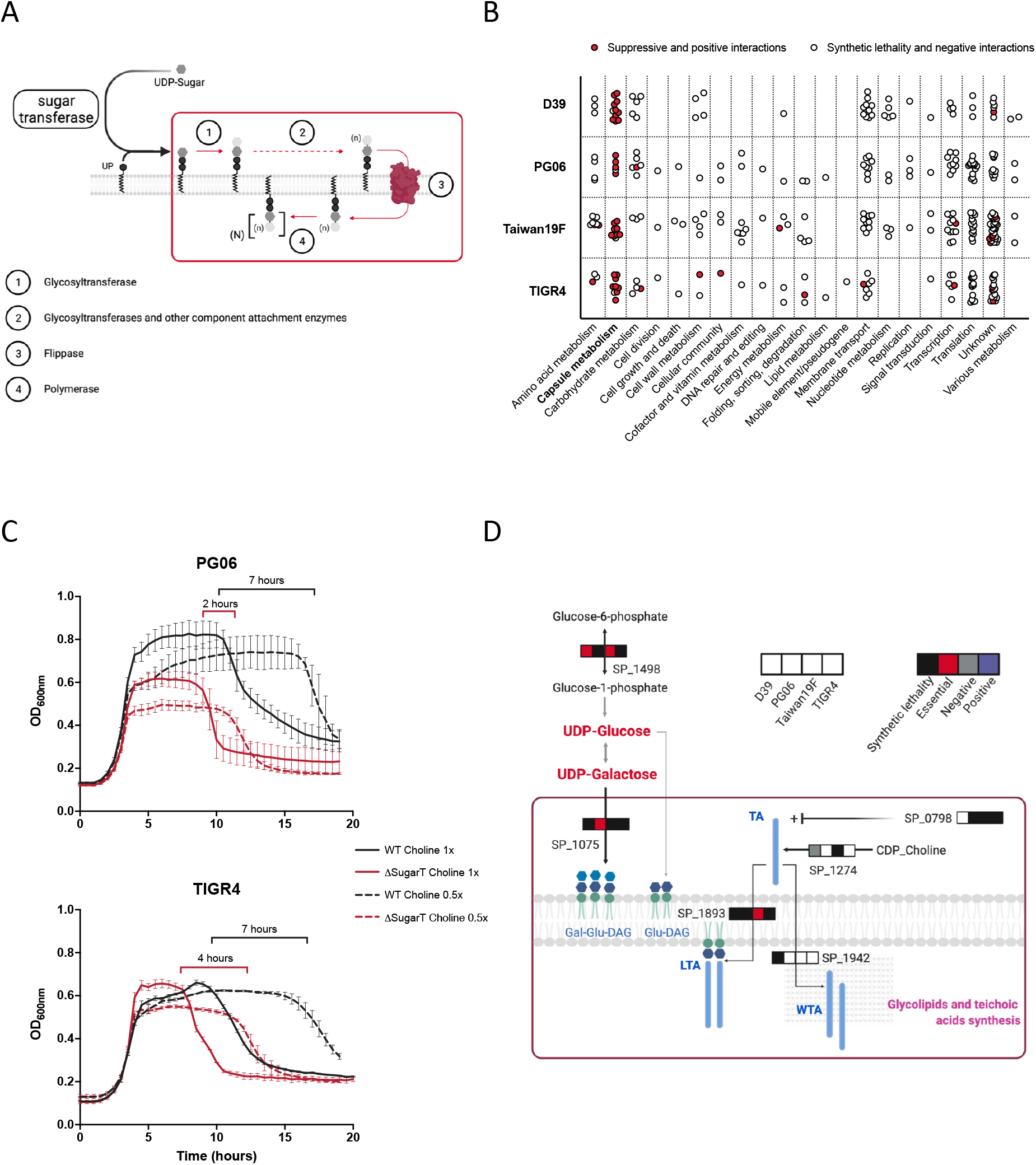
Capsule sugar transferase knockouts characterization. A) Schematic representation of capsule biosynthesis in *S. pneumoniae*. Genes inside the red box are accessory essentials. B) Genetic interactions identified by Tn-Seq using libraries constructed in sugar transferase knockouts derived from four different strains. Genes inside the red box in the right panel showed a suppressive interaction with the sugar transferase (red dots from Capsule metabolism category). C) Growth phenotype of wild type strains PG06 and TIGR4 and their derived knockouts in the capsule sugar transferase genes in medium SDMM with two different choline concentrations. The bars above curves indicate the time difference in the onset of autolysis between the two conditions for wild types (black bars) and knockouts (red bars). Data shown are the results from a single experiment with three replicates per condition (three different wells, same 96-well plate). Independent repetition of the experiment (>2) showed similar results. D) Genetic interactions identified by Tn-Seq using libraries constructed in sugar transferase knockouts derived from four different strains related to teichoic acid and glycolipids synthesis. The order of the squares in the strips indicates the strain and color the type of interaction identified. White square means no interaction. Red bold larger font indicates metabolites that accumulate in the context of *S. pneumoniae* non-encapsulated strains^81^. Gal-Glu-DAG: galactosyl-glucosyl-diacylglycerol, Glu-DAG: glucosyl-diacylglycerol, LTA: lipoteichoic acid, WTA: wall teichoic acid.

As previously described for unencapsulated strains of *S. pneumoniae*, the knockouts of the sugar transferase present an earlier onset of autolysis (Fig. 5C)^64, 65^. Autolysis in *S. pneumoniae* relies on the amount of LytA protein (encoded by SP_1937) bound to the choline moiety of wall teichoic acids^66^. Accordingly, the sugar transferase knockout-observed early autolysis relates to the amount of choline in the medium. Importantly, Tn-Seq in the background of the sugar transferase knockout reveals multiple synthetic lethal interactions with genes from a wide variety of functional categories including many involved directly and indirectly in cell wall and teichoic acid biosynthesis (Fig. 5B, D, and Extended Data Fig. 2). This suggests that teichoic acid production becomes more crucial when capsule synthesis is blocked. Moreover, the sugar transferase has a synthetic lethal interaction with the response regulator CiaR (SP_0798), which regulates teichoic acid biosynthesis genes^67–70^. The requirement of this transcriptional regulator and the observation that unencapsulated strains present more phosphorylcholine exposed on the cell surface^71^ suggest that teichoic acid production increases in the context of the sugar transferase knockout, and this increase may be the cause behind the early onset of autolysis of *S. pneumoniae* unencapsulated strains.

Overall, these data show that the essentiality of capsule biosynthesis genes can be bypassed by blocking the initiation of capsule synthesis. However, this interference with capsule synthesis comes at a cost, as indicated by the high number of synthetic lethal and negative interactions between the sugar transferases and other non-essential genes (Fig. 5B, Supplementary Table 8, and Extended Data Fig. 2). Importantly, this also highlights the potential for applying synergistic antimicrobials targeting accessory essentials and core genome genes.

### Essentiality of Magnesium transporter CorA relies on redundancy through a second transporter

Magnesium transporters described in bacteria belong to one of three classes: CorA transporters, MgtA/B, and MgtE transporters^72^. The *S. pneumoniae* core genome includes two members of the CorA family, SP_0185 and SP_1751 (Fig. 6A), with SP_0185 being essential in 8 out of 21 strains (Supplementary Table 4) and SP_1751 being non-essential. To determine why SP_0185 is differentially essential, we attempted to construct SP_0185 knockouts in 3 backgrounds the gene is essential in, and in 1 background (PG04) it is non-essential. We could only recover knockouts for PG04 (Fig. 3B), which suggests that strains where SP_0185 is essential cannot easily evolve mechanisms to bypass its essentiality. Tn-Seq genetic interaction screening using libraries constructed in the PG04 SP_0185 knockout shows that it has a synthetic lethal interaction with the non-essential transporter SP_1751 (Fig. 6B, Supplementary Table 8). The strain-dependent essentiality of SP_0185 therefore relies on potential functional redundancy with SP_1751.

**Figure 6:**
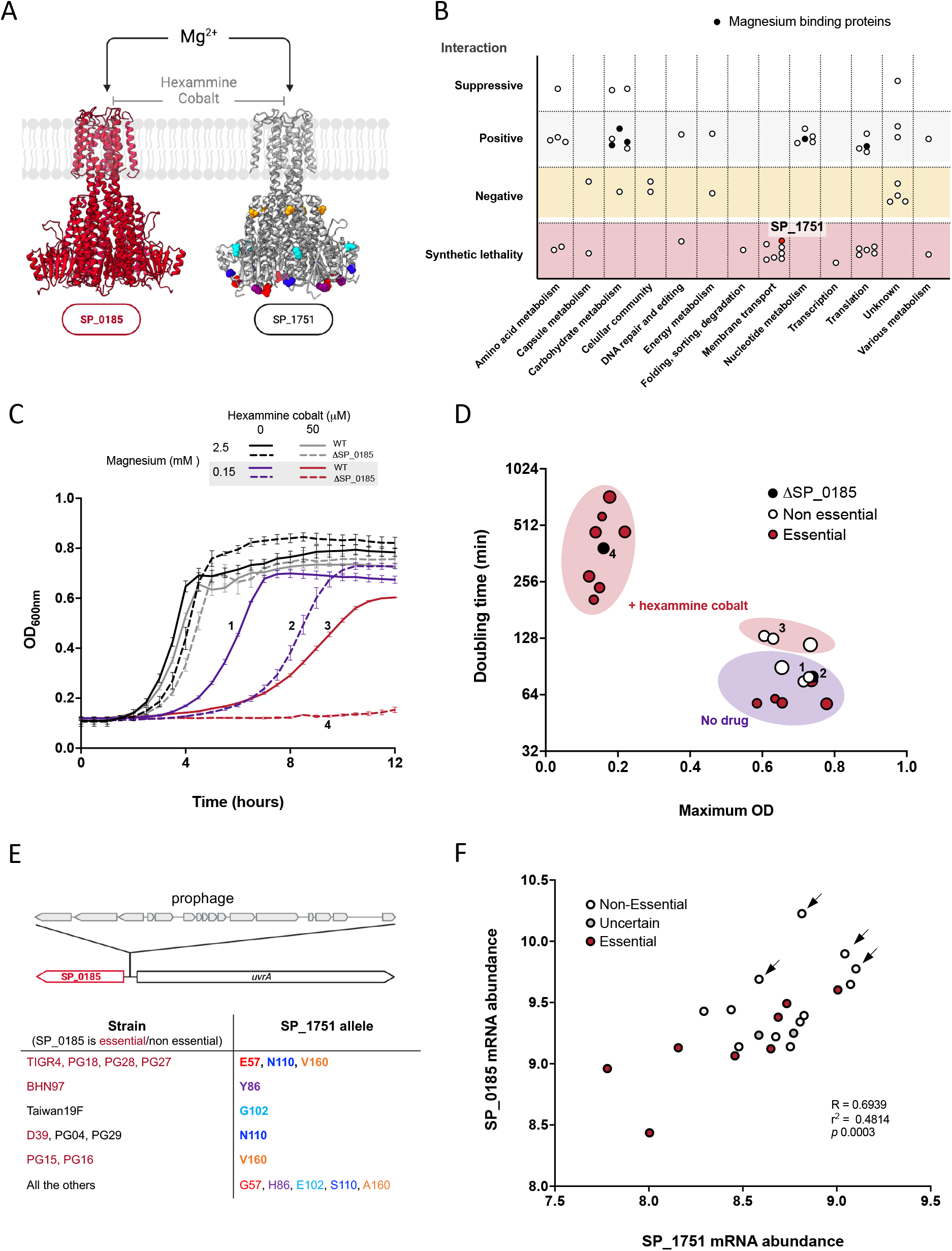
Redundancy in *S. pneumoniae* magnesium transport is strain-dependent. A) SWISS-MODEL predicted structures of the strain-dependent essential CorA magnesium transporter SP_0185 (Qmean = -2.27), and the second non-essential CorA transporter SP_1751 (Qmean = -2.82). Colors different than grey in the SP_1751 model show the residues with allelic variation across the PG collection strains. B) SP_0185 genetic interactions identified by Tn-Seq using libraries constructed in a PG04 SP_0185 knockout. Each dot represents one gene, and its graph location corresponds to the genetic interaction and functional category. Note that five of the 22 genes with positive interaction with SP_0185 are magnesium containing protein, suggesting that preventing the use of the metal for non-essential reactions can reduce the impact of the deletion. C) Growth of wild type strain PG04 (black and violet lines) and derived SP_0185 knockout (grey and red lines) in SDMM medium with different concentrations of MgCl_2_ and hexammine cobalt. Data shown are the results from a single experiment with three replicates per condition (three different wells, same 96-well plate). Independent repetition of the experiment (>3) showed similar results. D) Doubling time versus maximum OD observed for different strains growing in SDMM MgCl_2_ 0.15 mM in the absence or presence of the drug hexammine cobalt. Black dots represent the PG04 derived SP_0185 knockout. White and red dots represent wild type strains where SP_0185 is non-essential or essential, respectively. Dots numbered 1 to 4 correspond to the numbered curves in C. E) Different genomic neighborhoods of SP_0185 and allelic variation of SP_1751. The top drawing shows the most common genetic organization of SP_0185: next and in the opposite direction to the *uvrA* gene. Some strains have prophage sequences inserted between these two genes (grey ORFs above). The table at the bottom summarizes the different aminoacidic polymorphisms in the SP_1751 protein across the PG collection. Red font indicates those strains where SP_0185 is essential. Residues polymorphisms colors correspond to the SP_1751 model structure represented in A. F) Comparison of the transcript abundances of the two *corA* genes. Each dot represents a different wild type strain. Dot colors indicate if SP_0185 is essential or not. Arrows indicate those strains with the prophage inserted between SP_0185 and *uvrA*. R and *p* values derived from a least squares linear regression.

To confirm that the CorA transporters’ redundancy indeed lies in Mg^2+^ homeostasis, we evaluated the growth phenotype of nine wild type strains, and PG04 ΔSP_0185 at different Mg^2+^ concentrations and in the absence or presence of hexammine cobalt, a drug that specifically targets CorA proteins^73^ (Fig. 6C and D). Low Mg^2+^ concentration significantly affects the growth phenotype of ΔSP_0185 compared to the PG04 parental strain, while addition of the drug hexammine cobalt completely abolishes growth of the knockout (Fig. 6C). Furthermore, growth phenotypes of strains where SP_0185 is essential are similar to that of PG04 ΔSP_0185 (Fig. 6D). These results show that: 1) PG04 ΔSP_0185 acquires magnesium through another transporter sensitive to hexammine cobalt, such as SP_1751, corroborating the identified synthetic lethality between the two transporters; and 2) the strains where SP_0185 is essential rely predominantly on one transporter to acquire the ion, and thus SP_1751 is not necessarily redundant with SP_0185 in the context of these strains.

To shed more light on this strain-dependent redundancy, we explored these two genes further with respect to their genomic context, protein sequence, and expression level (Fig. 6E and F). First, two genomic arrangements exist for gene SP_0185 across the PG collection (Fig. 6E top panel). In the most common configuration, the transporter is next to *uvrA,* and oriented in the opposite direction of this gene. In the second configuration a prophage sequence is present in between both genes (Fig. 6E). Second, while the SP_0185 protein sequence is identical across strains, the SP_1751 transporter exhibits polymorphisms. For instance, E57, Y86 and V160 are polymorphisms in SP_1751 that appear only in strains where SP_0185 is essential (Fig. 6E bottom panel). Third, expression levels of the transporters are positively correlated (Fig. 6F). In general, strains with the prophage inserted next to SP_0185 display some of the highest expression levels for both transporters and are the strains in which SP_0185 is non-essential (arrows in Fig 6F). In contrast, the backgrounds in which SP_0185 is essential mostly display a lower expression level for both transporters. Overall this suggests that a high expression of SP_1751 in combination with several specific polymorphisms in this gene can overcome the absence of SP_0185. However, while the magnesium transporters are at least partially redundant in a strain dependent manner, on the species level SP_0185 is the main magnesium transporter.

### S. pneumoniae easily evolves mechanisms to overcome farnesyl-PP-synthase essentiality

Undecaprenyl phosphate (UP) is a crucial bacterial molecule involved in translocating peptidoglycan, teichoic acids, and capsule intermediates from the cytoplasm to the outer leaf of the membrane^74–76^. In the UP-biosynthesis pathway, the enzyme encoded by the gene SP_1205 catalyzes the condensation between isopentenyl diphosphate (IPP) and geranyl diphosphate (GPP) into farnesyl diphosphate (FPP), which is then converted to undecaprenyl diphosphate (UPP) by the enzyme encoded by SP_0261. Lastly, UPP is dephosphorylated into its active form UP by the SP_0457 encoded enzyme (Fig. 7A). In our analysis, SP_0261 is a universal essential gene, while SP_1205 is a strain-dependent essential gene (Fig. 7A, Supplementary Table 5).

**Figure 7:**
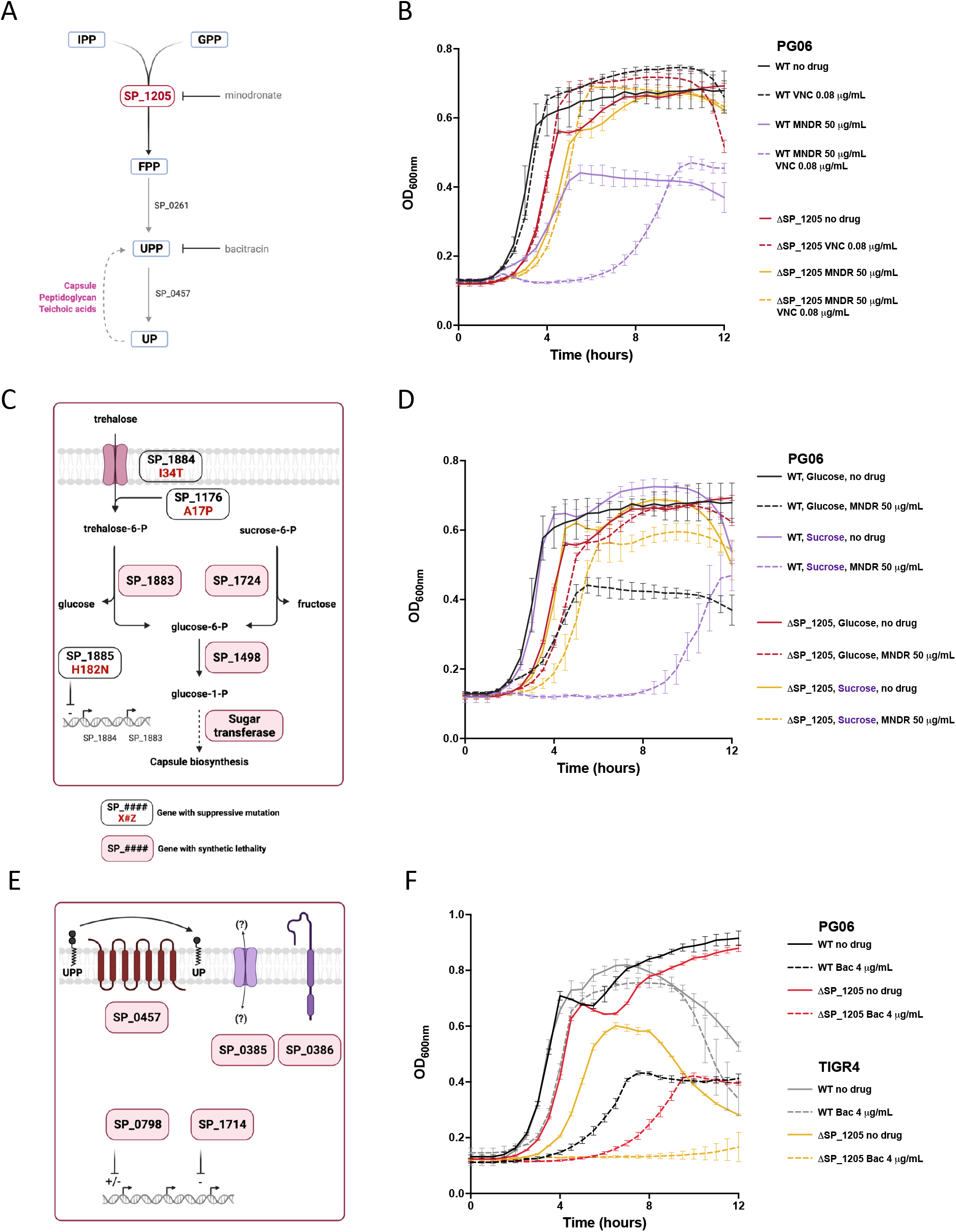
Farnesyl-PP synthase (SP_1205) essentiality is easily bypassed. A) Role of the product of the strain-dependent essential gene SP_1205 in the synthesis pathway of undecaprenyl phosphate (UP). B) Synergism between minodronate (MNDR) and vancomycin. Growth phenotype of wild type strains PG06 (SP_1205 is non-essential) and the derived SP_1205 knockout in medium SDMM in the presence of non-inhibitory concentrations of vancomycin (VNC), in the presence or absence of the SP_1205 targeting drug minodronate (MNDR), or both. C) Disaccharide metabolism related genes probably involved in mechanisms of SP_1205 essentiality bypass. Selection of suppressive mutations of SP_1205 essentiality identified by *breseq* (boxes with white background) and synthetic lethality interactions (boxes with pink backgrounds) identified by Tn-Seq using libraries constructed in a PG06 SP_1205 knockout. D) Carbon source and SP_1205 requirement. Growth phenotype of wild type strains PG06 (SP_1205 is non-essential) and the derived SP_1205 knockout in medium SDMM with glucose or sucrose as carbon source, and with or without MNDR. E) Bacitracin resistance related genes probably involved in mechanisms of SP_1205 essentiality bypass. Selection of synthetic lethality interactions (boxes with pink backgrounds) identified by Tn-Seq using libraries constructed in a PG06 SP_1205 knockout. F) SP_1205 knockouts bacitracin sensitivity. Growth phenotype of wild type strains PG06 (SP_1205 is non-essential), TIGR4 (SP_1205 is essential), and their derived SP_1205 knockouts in medium SDMM in the presence or absence of the UPP recycling inhibitory drug bacitracin (Bac). Data from B, D, and F are the results from single experiments with three replicates per condition (three different wells, same 96-well plate). Independent repetition of the experiment (>3) showed similar results. FPP: (2*E*, 6*E*) farnesyl diphosphate, GPP: geranyl diphosphate, IPP: isopentenyl diphosphate, UP: undecaprenyl phosphate, UPP: undecaprenyl diphosphate.

Transformation experiments targeting SP_1205 confirmed the non-essentiality of the gene in the PG06 background (Fig. 3B). In contrast, transformation in strains PG16 and TIGR4 (in which the gene is essential) yielded very few colonies. Excitingly, while WGS confirmed that these colonies lack a functional copy of SP_1205, multiple putative suppressive SNPs (Supplementary Table 7) were present in these colonies, suggesting that *S. pneumoniae* may be able to evolve different mechanisms to bypass SP_1205 essentiality.

To gain further functional insights into SP_1205, we evaluated growth of different strains in the presence and absence of the drug minodronate (MNDR) (Fig. 7B, Extended Data Fig. 3A). MNDR is a bisphosphonate drug currently in a phase-3 trial for human use, which targets human FPP synthase to suppress bone resorption and bone loss. However, there is evidence that bisphosphonates may also have antimicrobial potential^77–79^. Indeed, the drug inhibits growth of several wild-type strains. In contrast, in strains where SP_1205 is missing because it is non-essential (PG06) or because of bypass mutations (PG16, TIGR4) growth is unaffected. Additionally, because UP is critical for the cell wall, inhibition of UP synthesis may thus synergize with cell-wall synthesis inhibitors, like vancomycin^80^. Accordingly, the combination of MNDR with vancomycin at a subinhibitory concentration has a synergistic effect on a wild type strain’s growth (Fig. 7B, Extended Data Fig. 3A), while there is no effect in ΔSP_1205 strains. These results confirm that MNDR specifically targets the SP_1205 gene-product.

WGS and Tn-Seq in a ΔSP_1205-PG06 query strain revealed that many of the putative bypass mutations and synthetic lethal interactions associate in two different cellular processes: **1)** the utilization of disaccharides leading to capsule biosynthesis; and **2)** the recycling of UP. For the first process, the identification of three functionally related suppressive SNPs (SP_1176 A17P; SP_1884 I34T; SP_1885 H182N) suggests that transport of the disaccharide trehalose (or a similar compound) is a central hub for evolving SP_1205 bypass mechanisms (Fig. 7C). Gene SP_1205 has a synthetic lethal interaction with two enzymes (SP_1883 and SP_1724) that hydrolyze the disaccharides trehalose and sucrose to produce glucose-6-phosphate, which feeds into glycolysis, the pentose-phosphate pathway, and capsule biosynthesis. Based on these results, we evaluated the effect on growth with the disaccharide sucrose as the carbon source (Fig. 7D, Extended Data Fig. 3B). For wild type strains, sensitivity to MNDR increases when growing with sucrose, showing that SP_1205 becomes more crucial when this disaccharide is the main carbon source. Tn-Seq experiments using both PG06 sugar transferase and SP_1205 knockout libraries show that the sugar transferase has a robust negative interaction with SP_1205, demonstrating that capsule production is essential to growth when SP_1205 is deleted (Fig. 7C, Supplementary table 8). Additionally, it has been shown that growth in the presence of sucrose negatively affects *S. pneumoniae* capsule production^81^. Transposon insertions in the hydrolases genes SP_1883 and SP_1724 may thus inefficiently process disaccharides and thereby trigger a lower capsule production, explaining their synthetic lethality with SP_1205.

The second group of suppressive SNPs and genetic interactions relate to UP recycling (Fig. 7E, Supplementary Tables 7 and 8). UP homeostasis depends on a critical balance between demand, synthesis and recycling^76^, and SP_1205 sits at the synthesis root of this balance. Bacitracin is a drug that binds to UPP, preventing its recycling to UP^82^. Importantly, we identified different genes previously described as induced by and required to overcome bacitracin bactericidal effect as having synthetic lethality with SP_1205 (Fig. 7E, Supplementary Table 8)^82–84^. One of these genes with synthetic lethality is the UPP phosphatase BacA (SP_0457). Interestingly, a TIGR4 derived SP_1205 knockout presented two compensatory mutations associated with the cell wall: one mutation is located in *murE* (SP_1530 A430S), which is involved in peptidoglycan synthesis, and the second mutation is located in a non-characterized membrane protein coding gene (SP_0454 H581N), which forms a complex with BacA^85^. We thus hypothesized, that a ΔSP_1205 strain would be more sensitive to bacitracin. Indeed, sensitivity to bacitracin is much higher in ΔSP_1205 strains, confirming that when synthesis is inhibited, recycling of UPP becomes more crucial (Fig. 7F, Extended Data Fig. 3C). Interestingly, PG06-WT, in which SP_1205 is non-essential, is more sensitive to bacitracin than for instance TIGR4 (in which SP_1205 is essential), suggesting that in PG06 UP homeostasis is at default more skewed towards recycling. Importantly, these data show that *S. pneumoniae* can evolve different solutions to make SP_1205 dispensable. However, besides these solutions, any stress that affects the homeostasis of UP (bacitracin) or capsule production (sugar transferase deletion, accumulation of disaccharides) makes SP_1205 essential.

### Formate tetrahydrofolate ligase essentiality relates to the stringent response

The enzyme encoded by SP_1229 belongs to the One Carbon pool by folate pathway (KEGG:00670). In this pathway, tetrahydrofolate derivatives serve as carbon donors for different reactions, like the synthesis of purines, thymidylate, methionine, serine, and glycine (Fig. 8A)^86, 87^. We successfully recovered knockout mutants for two strains where the gene is non-essential (PG04 and TIGR4). Additionally, knockouts were obtained for PG16, in which the gene is essential, but only in combination with different SNPs, likely acting as suppressors (Fig. 8A, Supplementary Table 7). Two of these mutations, present in two independent backgrounds, create premature stop codons in SP_0918 and SP_1378, which connect to the One Carbon pool by folate pathway through the S-adenosyl-methionine cycle. SP_0918, when deleted in strain D39, diminishes the production of spermidine^88^, while SP_1378 codes for a SAM-dependent RNA-methyltransferase. Moreover, three other PG16 SP_1229 knockouts showed additional suppressor mutations in genes involved in guanosine ribonucleotides metabolism and (p)ppGpp biosynthesis (SP_0012 T104A and SP_1645 S310F, SP_1645 A378T, and SP_1738 Y35C). As above, these results indicate that mutations in associated pathways can help overcome essentiality, in this case for SP_1229.

**Figure 8:**
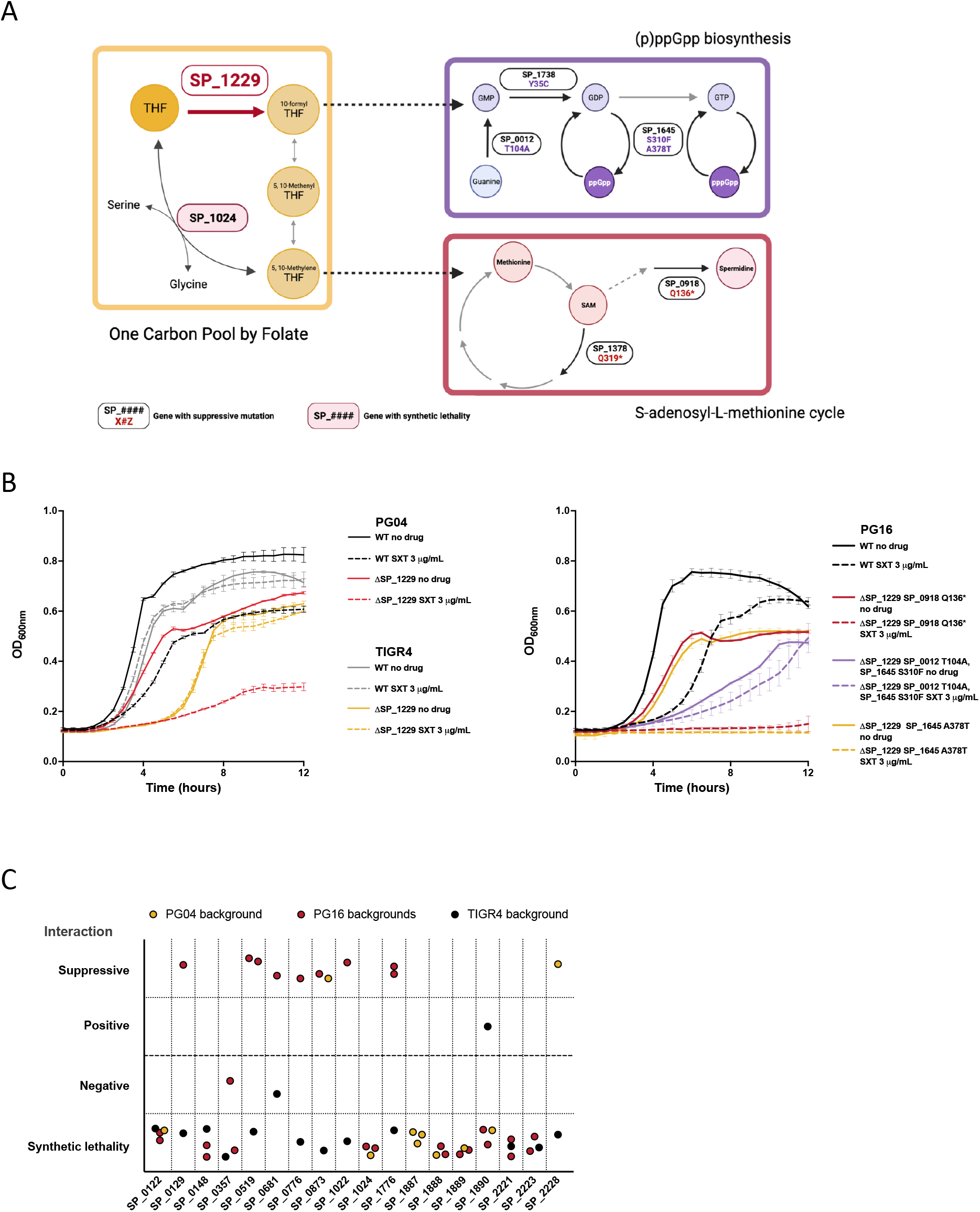
*S. pneumoniae* strains have different mechanisms to counteract formate tetrahydrofolate reductase (SP_1229) deletion. A) Role of the product of the strain-dependent essential gene SP_1229 in the One Carbon Pool by Folate pathway and suppressive mutations of SP_1229 essentiality identified by *breseq* (boxes with white background) in different PG16 strain derived knockouts. Dashed arrows indicate multiple non-shown metabolic reactions that connect represented pathways. B) Growth phenotype of wild type strains PG04, TIGR4, PG16, and their derived SP_1229 knockouts in medium SDMM in the presence or absence of the THF synthesis inhibitory drug cotrimoxazole (SXT). SP_1229 is non-essential in PG04 and TIGR4, and essential in PG16 background. Data are the results from single experiments with three replicates per condition (three different wells, same 96-well plate). Independent repetition of the experiment (>3) showed similar results. C) Selection of genetic interactions identified by Tn-Seq using libraries constructed in two backgrounds where SP_1229 is non-essential (PG04 and TIGR4) and two knockouts derived from PG16 with different suppressive mutations (SP_0918 Q136* and SP_1378 Q319*). SAM: S-adenosyl-L-methionine, THF: tetrahydrofolate.

To investigate how strains may bypass SP_1229 essentiality in different ways, we evaluated growth in the presence or absence of the drug cotrimoxazole (SXT) (Fig. 8B). This drug inhibits the synthesis of the SP_1229 substrate tetrahydrofolate. PG04 and the PG16 derived knockouts grow slower in rich medium compared to WT, and except for one of the PG16 derived knockouts have increased sensitivity to SXT. In contrast, the TIGR4 derived knockout grows similar to the wild type. These results show that besides the strain-dependent essentiality of SP_1229, the effects of deleting the gene differ across strains where the gene is non-essential.

Tn-Seq experiments using libraries constructed in the PG04, TIGR4, and two different PG16 derived knockouts revealed shared and strain-dependent genetic interactions with SP_1229 (Supplementary Table 8). Interestingly, nine genes have an opposite genetic interaction in TIGR4 compared to the other three backgrounds (Fig. 8C). A key interaction exists with the serine hydroxymethyltransferase gene SP_1024, which shows synthetic lethality with SP_1229 in PG04 and PG16, but not in TIGR4. This enzyme can bypass SP_1229 by using serine as substrate (Fig. 8A)^89^. Two lines of evidence suggest that this SP_1024-dependent bypass mechanism links with (p)ppGpp metabolism. First, some strains like PG16 require additional changes in associated pathways in order to bypass SP_1229 essentiality, including suppressor SNPs in the (p)ppGpp synthetase enzyme RelA (SP_1645), and two related enzymes: the guanine phosphoribosyltransferase SP_0012, and the guanylate kinase SP_1738. Second, in strain D39, mupirocin, a drug that robustly triggers the stringent response induces the expression of SP_1024 in a SP_1645 dependent manner^90^. Overall, these results show that different bypass mechanisms exist to overcome SP_1229 inactivation.

## Discussion

In this work we use a systems biology approach to determine, at the species level, how the bacterial pan-genome might influence gene-essentiality, and whether essential genes that are initially critical for the survival of an organism can evolve to become non-essential. Our strategy identified a set of 206 universally essential genes whose essentiality cannot be circumvented by the genetic background. Additionally, we identified 186 strain-dependent essential genes for whom the genetic background is a significant modulator of essentiality. Consequently, we demonstrate that different *S. pneumoniae* strains were able to evolve mechanisms to bypass some of these gene’s essentiality. An experimental advantage of this strain-dependent essentiality is that it enables an approach to untangle gene function in backgrounds where a gene is non-essential. Additionally, for those that have a high evolvability potential, gene function can be unpacked in strains that can acquire compensatory mechanisms. Taking advantage of this we revealed novel aspects of *S. pneumoniae* biology and defined mechanisms behind multiple strain-dependent genes, including: **1)** modulation of the composition of the polar lipid membrane (SP_1603); **2)** the presence/absence of accessory genes (SP_1809/SP_1569); **3)** prevention of toxic stalling of the capsule biosynthesis pathway; **4)** the ability to use redundant genes (SP_0185); **5)**a fine-tuned balance between synthesis, demand and recycling of crucial cell-wall components (SP_1205); and **6)** The ability to rewire pathways in order to fulfill synthesis demands of specific metabolites (SP_1229).

Analogous strategies could be implemented for other human pathogens with well described pan-genomes by employing Tn-Seq or similar tools (e.g., CRIPSRi), in order to shed more light on the fundamentals of gene essentiality and to delineate the evolvability-potential of essential genes. Importantly, *S. pneumoniae* shares at least two key characteristics with many other bacterial pathogens: **1)** the species is represented by a large pan-genome; and **2)** the emergence of antibiotic resistant and tolerant strains compromises the future success of current therapies. Therefore, we expect that our main findings extend beyond *S. pneumoniae* biology, with at least two important consequences. First, our work suggests that gene essentiality may be less rigid than currently assumed. Accordingly, rather than classification of genes as essential, it may be more useful to categorize essential genes in a more quantitative manner^7^ that captures how context-dependent essentiality is, and how likely a gene is to remain essential. The second consequence is related to essential genes as targets for the development of new therapeutics. Our findings that many essential genes are strongly influenced by the genetic-background, and thereby may have a high evolvability-potential, suggests that this could undermine the attractiveness of strain-dependent essentials as candidate drug targets. However, we also show that elimination of strain-specific essentials come with a large fitness cost *in vitro*, and especially *in vivo,* indicating that drugs inhibiting the products of strain-dependent essentials would still be effective in targeting a bacterial pathogen. Moreover, our genetic interaction studies show that the majority of the synthetic lethality interactions identified here exist between strain-dependent essentials. This observation combined with our drug-sensitivity assays, exemplified by: **a)** MNDR, a repurposed drug that targets a strain-dependent essential gene and synergizes with cell-wall synthesis inhibitors; and **b)** drugs such as SXT that block folate synthesis, which synergize with drugs inhibiting enzymes of the one-carbon pool by folate pathway, highlight that strain-dependent essentials could also be promising targets of synergistic therapies.

Lastly, a pan-genome with strain dependent essentiality, and more specifically, strains in which essentiality can be altered by changes to the genetic background including bypass mutations, may also create new targeted drug screening opportunities. For instance, the sugar transferase knockout strain that bypasses essentiality of capsule biosynthesis can be screened in parallel with the wildtype strain for drugs that target capsule biosynthesis for which the wildtype would be sensitive, but the mutant would not. Different sets of strains with changes to different pathways and/or processes which thereby enable differential essentiality could thus be used to identify compounds that target a specific gene-product and/or process. We are currently exploring these and other approaches, which highlights how exploration of basic biology can potentially go hand in hand with the development of new antimicrobial strategies.

## Methods

### Strains and growth conditions

*S. pneumoniae* strains used in this study are listed in Supplementary Table 1. Thirty of them (PG01 to PG30) belong to a surveillance study done in a Nijmegen hospital, NL^25^, while the other six are lab strains available at our lab.

Different strains were grown on Tryptic Soy Agar or Blood Agar Base no. 2 (Sigma-Aldrich) plates supplemented with 5% defibrinated sheep’s blood at 37°C in a 5% CO_2_ atmosphere. Liquid cultures were grown statically in THY, C+Y or semi-defined minimal media (SDMM) at pH 7.3, with 5 µl/ml Oxyrase (Oxyrase, Inc)^2^ at the same incubation conditions as plates. For growth curve assays, strains were grown in THY until an OD_600_ of ∼0.5, pelleted and resuspended in same volume of phosphate saline buffer (PBS). OD_600_ was adjusted to 0.05 in PBS and 20 μL of this suspension were diluted in 180 μL different media conditions in wells of flat bottom 96-well plates. OD_600_ measurements were taken on a BioSpa 8 plate reader (BioTek), and experiments were repeated at least three times. Different modifications to SDMM were assayed as described in figure legends and text.

### Genomic DNA isolation and PacBio sequencing

To obtain high-quality and high-yield of genomic DNA for PacBio sequencing and Tn-Seq libraries construction, strains were grown in 20 mL of THY (plus oxyrase, plus catalase 200 units/mL) until an OD_600_ of ∼0.7, and cells were recovered by centrifugation (3000g, 7 min). Pellets were subsequently processed and gDNA purified using QIAGEN Genomic-tip 100/G columns. Initial lysis of cells was achieved by resuspending pellets in a lysis buffer adapted for *S. pneumoniae* (10 mM Tris·Cl, pH 8.0; 10 mM EDTA pH 8.0; 0.1% Tween®-20; 1.1% Triton X-100, sodium deoxycholate 1.5%) containing 200 μg/mL of RNAse A (Macherey-Nagel), and incubated during 15 min at 37 °C. The rest of the protocol was performed as described in the QIAGEN genomic DNA handbook. For mucoid strains, like PG23 and PG24 (serotype 3), cells were washed once with NaCl 1M, and once with PBS before proceeding with lysis.

Single-molecule real-time sequencing (SMRT-Seq) was carried out on a PacBio Sequel-I instrument (Pacific Biosciences, USA). Genomic DNA concentration was determined using the QuBit dSDNA HS (High Sensitivity) assay kit (Thermo Fisher), and purity was calculated using a Nanodrop Spectrophotometer 1000. Genomic DNA samples (3 μg) were sheared to an average size of 10 kb via G-tube (Covaris, Woburn, MA) before library preparation. Libraries were then generated with SMRTbell Express Template Prep Kit 1.0 and pooled libraries were size selected using the BluePippin system with 0.75% Pippin Gel Cassettes and Marker S1 (Sage Sciences) at a 4kb minimum threshold. Sequencing reads were processed using the Pacific Biosciences’ SMRTAnalysis pipeline version 8.0.0.80529 and assembled using Microbial Assembler.

Closed single-contigs were annotated using the NCBI Prokaryotic Genome Annotation Pipeline (PGAP) and genome assemblies and assembly information is made available as part of the GenBank Bioproject PRJNA514780.

### Pan-genome core, accessory genome, and phylogeny determination

Three pipelines for orthologous gene clustering and pan-genome analysis (PanX^31^, PPanGGOLin^32^, BF-Clust^33^) were applied to a study group of 208 sequenced isolates including the 36 strains of our collection (Supplementary Table 2). In the case of BF-Clust methodology, clusters were subclustered according to their genomic neighborhood in order to discriminate between paralogs (Supplementary Table 3). When referring to specific genes, we use as a convention the old locus tag from TIGR4 (SP_#). For those genes not present in TIGR4 genome we will use D39 locus tag (SPD_#), or Taiwan19F (SPT_#), or the BF-Clust subcluster ID (#_#).

Phylogenetic tree of figure 1B was obtained as an output from the PanX pipeline, constructed based on the core genes SNPs of the selected 208 strains, and edited using iTol^91^.

### Tn-Seq libraries construction

Six independent transposon libraries, each containing 10,000 to 20,000 insertion mutants, were constructed with transposon Magellan6 in the different strains as described previously^19, 41, 42, 92^, with the following modifications: 1) gDNA used was isolated as described for PacBio sequencing, 2) Marc9 transposase expression was induced with IPTG 0.5 mM in medium 2YT with ethanol 2%, at 25°C during 4 hours, 3) Transformation reactions were scaled up to 4 mL volume. Transposon mutants were recovered in Blood Agar Base no. 2 plates supplemented with 5% sheep’s blood with 200 μg/mL spectinomycin. Libraries stock cultures were grown several independent times in THY medium, genomic DNA was isolated using Nucleospin tissue kit (Macherey-Nagel), and Tn-Seq sample preparation, and Illumina sequencing were done as described^12, 19, 41, 42, 92, 93^.

### Tn-Seq essential genes, fitness and genetic interactions determination

Binomial method of the TRANSIT package was used for determination of essential genes in the 22 strains with constructed libraries^37, 38^. Binomial categorization relies on the calculation of a z-value determined by library saturation, the number of TA sites in the gene, and a calculated FDR of 5% to set the “Non-essential” and “Essential” z-value thresholds. “Uncertain” predicted genes have z-values between these thresholds. 10% of the sequence of a gene from the 5’ and 3’ ends were omitted for the calculation of the z-value. To compare between strains and determine the essential genes classes, only strains with a saturation higher than 35% were considered (17 strains). The maximum and minimum z-values observed for each gene across strains were calculated. All genes with a maximum z-value above the essential threshold were included as part of the essentialome. Core genes with minimum z-value above the essential threshold were classified as Universal essentials. Core genes with maximum z-values above the essential threshold (then part of the essentialome), but minimum z-values below the essential threshold were classified as strain-dependent essential genes. One caveat to consider about this last classification is that for some genes minimum z-values are in the Uncertain zone (between essential and nonessential thresholds). This could lead to some false positives, especially for those genes with minimum z-values close to the essential threshold.

To obtain the average mutant fitness (W̄) in medium SDMM, Tn-Seq libraries were first grown in THY until an OD_600_ of ∼0.5, pelleted and diluted to an OD_600_ of 0.1 in SDMM with 20 mM glucose as carbon source. Cells were let grown for 4 generations (∼3 hours). Genomic DNA from THY and SDMM grown cells were processed for Tn-Seq. W_i_ (where I denotes a specific insertion mutant) for each transposon insertion is calculated as previously described by comparing the fold expansion of the mutant relative to the rest of the population at the initial time (THY) and the final time (SDMM)^41, 42, 93^. All of the mutants W_i_ in a specific gene, excluding those with insertions in the first and last 10% of the gene total sequence, are then used to calculate the W̄ and standard deviation for the gene in question. To determine statistically if a gene interruption affects growth in SDMM the following analysis for each strain is performed: 1) Essential genes specific for each strain and genes with less than three data points are excluded from the analysis, 2) median average fitness across all genes is calculated (based on results from Tn-Seq experiments with high bottleneck, we assume that the median is the better estimate for a fitness value that represents no effect on growth), 3) the observed W̄ and the median (expected W) are compared by a one sample *t*-test, and obtained p-values are corrected for multiple comparisons by a FDR of 5% using the two-stage step-up method of Benjamini, Krieger and Yekutieli ^94^, 4) genes tagged as a true discovery and with a ΔW (W̄ observed – W expected, median) absolute value higher than 0.2 are considered required (ΔW<-0.2) or disadvantageous (ΔW>0.2) for growth in SDMM.

Genetic interactions were determined by comparing Tn-Seq Binomial predictions (suppressive and synthetic lethality interactions) or SDMM fitness calculations (negative and positive interactions) between libraries constructed in wild type backgrounds and specific knockout backgrounds. Genes that changed their categorization from “Essential” in the WT context to “Non-essential” in the knockout context represent suppressive interactions, while genes that change the opposite way represent synthetic lethality. To reduce false negatives, genes that changed their categorization from “Uncertain” in WT to “Essential” in knockout backgrounds were tagged also as synthetic lethality. These last synthetic lethality interactions have then a lower confidence, especially for those “Uncertain” genes with binomial z-values closer to the “Essential” threshold. Positive and negative interactions are determined as follows: 1) SDMM growth rate ratio between site-directed constructed knockouts and WT was used as multiplicative model correction factor, 2) each mutant W̄ obtained from libraries constructed in the knockout and WT were multiplied by the correction factor, 3) the corrected W̄ s are compared by a two sample *t*-test, and obtained p-values are corrected for multiple comparisons by a FDR of 5% using the two-stage step-up method of Benjamini and Hochberg^95^, 4) genes tagged as a true discovery and with a ΔW (W̄ knockout corrected – W̄ wild type corrected) absolute value higher than 0.2 are considered to have a negative (ΔW<-0.2) or positive (ΔW>0.2) interaction with the respective knocked out gene.

### Transcriptome

Cells were grown in medium THY until an OD_600_ of ∼0.3 in triplicate. Then, RNA isolation, sample preparation, and Illumina sequencing were performed as previously described^20^. Raw data analysis process was also performed as described in Zhu et al., 2020, with the exception that aggregated feature counts were TPM normalized to compare across the different genes ^96^.

### Site-directed gene deletions and whole-genome sequence analysis

Site-directed gene knockouts were constructed by replacing target gene sequences with a chloramphenicol and/or spectinomycin resistance cassette as described previously^41, 42, 97^, but using up to 1000 ng of the allele exchange PCR product. All PCR reactions were done using Q5 polymerase (NEB). Primers used and knockouts constructed are described in Supplementary Tables 9 and 10. Randomly selected colonies and in some cases, whole transformation populations were recovered and their genomic DNA was isolated using Nucleospin tissue kit (Macherey-Nagel). Deletions were confirmed by two PCR reactions using primers outside of the PCR product and primers specific to the antibiotic marker used (F0-MR, R0-MF, Fig. 3A). Products of around 2kb are expected for the two PCR reactions. The amplification of a product of around 4kb in one of the reactions indicates the clone or population consist of merodiploids.

Complementation of SP_1603 knockouts was achieved by cloning WT copy of SP_1603 gene under the control of the constitutive promoter P3 at the non-coding region downstream SP_1885 (CEP locus)^23^.

For identification of putative suppressive mutations, strains genomic DNA concentrations were measured on a Qubit 3.0 fluorometer (Invitrogen) and diluted to 10ng/uL in Tris HCl 5mM for library preparation using a Nextera kit (Illumina). Libraries were sequenced on an Illumina NextSeq500 and reads were mapped to their corresponding reference genomes. Single nucleotide mutations, deletions, and new arrangements were identified using the breseq pipeline^98–100^. Merodiploids were distinguished from knockouts by PCR as described above or breseq. To distinguish merodiploids from knockouts, sequencing coverage of the target gene and the homology arms in the 3Kb product (Fig. 3C) were examined. In a merodiploid, coverage of left and right arms is about the double of the average coverage due to duplication, while the coverage of the target gene remains the same as the average. Another characteristic of merodiploids identifiable by *breseq* is that the left and right homology arms are jointed with a frequency around 50%.

Some suppressive mutations were validated as follow: 1) WT strains were co-transformed with a spectinomycin cassette that inserts in the neutral region between genes SP_2105-SP_2106 and a PCR product with the SNPs to validate in a 1:10 ratio. It is expected that a 20% of spectinomycin resistant colonies obtained should contain the SNP^101^. Whole population was pooled and transformed with the target essential suppressed gene.

### Membrane polar lipid composition determination

Strains were grown in C+Y medium at 37 °C to an OD_600_ of 0.2, and 50 μCi of [1-^14^C]acetate was added. After a two-hour labeling period, cells were harvested and washed twice with PBS. The lipids were extracted by the Bligh and Dyer method ^102^, and total incorporation was quantified by scintillation counting using Tri-Carb 2910 TR (Perkin Elmer). The lipids were separated by loading equivalent amounts of radioactivity on a Silica Gel 60A thin-layer plates (Partisil LK6D; Whatman) and developed in chloroform:methanol:acetic acid:water (80:10:14:3, v/v/v/v). After drying, the thin layer plate, was exposed to a PhosphorImaging screen overnight. The signal intensity was read using a Typhoon FLA 9500 (GE Healthcare Life Sciences) PhospohoImager and the distribution of label quantified using ImageQuant TL (GE Healthcare Life Sciences). The results are from two independent experiments.

### Data visualization and statistics

Figure panels were created by Biorender.com, and GraphPad Prism 9. CorA protein structures were modelled by SWISS-Model^103^ and edited using Chimera.X.1.2.5. Statistics analysis were performed using GraphPad 9 and R 1.4.1106.

### Data availability

Genome assemblies, assembly information, Tn-Seq, RNA-Seq and WGS raw data are available as part of the Bioproject PRJNA514780 on GenBank and SRA.

## Acknowledgements

DNA sequencing was performed at the Boston College Sequencing Core. The authors wish to thank the invaluable contributions of Amelieke Cremers, for sharing the strains isolated in Nijmegen, Jelena Radivojac and Sean Cotton for their help in RNAseq and PacBio sequencing experiments. Authors wish to thank also Zeyu Zhu, Bharathi Sundaresh, and Suyen Espinoza for valuable discussions. This work was supported by a PEW Latin America Fellowship and Charles King Trust Fellowship to F.R., NIH National Institute of Allergy and Infectious Diseases (grant R01 AI110724 to T.v.O., and grant U01 AI124302 to T.v.O. and J.W.R.), and through the NIH National Institute of Dental and Craniofacial Research (grant R01DE027850 to C.D.J.).

## Author contributions

F.R and T.v.O. devised the study and wrote the manuscript. F.R., E.R., J.L., and M.F. performed wet-lab experiments and data collection. D.S., J.L., and D.S.J. performed pan-genome ortholog cluster analysis. J.A. and D.S. contributed to Tn-Seq dry-lab analysis pipeline. J.W.R., and C.R. contributed to key conceptual ideas. C.D.J. performed PacBio sequencing analysis. F.R and T.v.O performed data analysis and interpretation. All authors contributed to manuscript editing and approved the final paper.

---

**Extended Data Figure 1:**
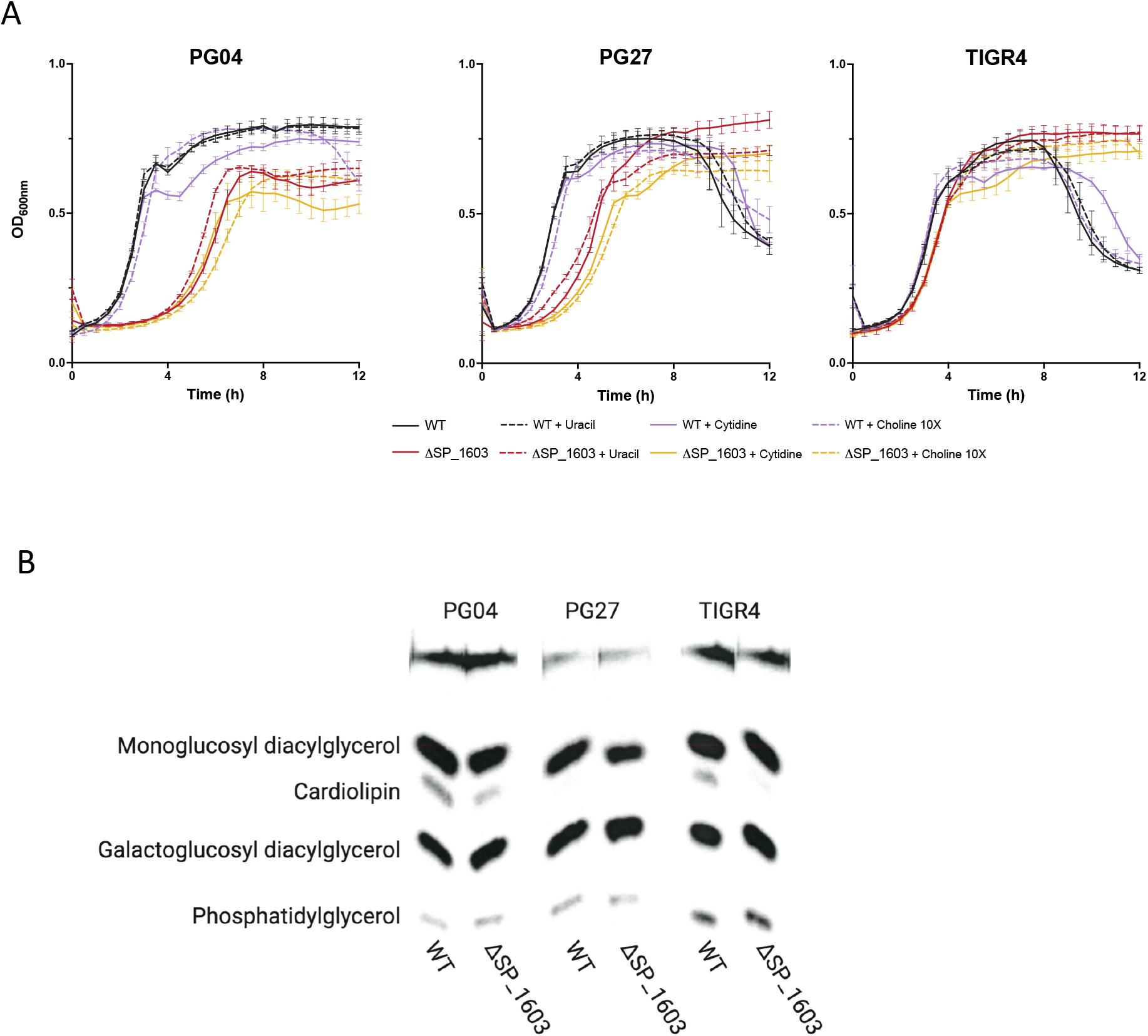
SP_1603 knockouts characterization. A) Growth phenotype of wild type and derived SP_1603 knockouts in medium SDMM with uracil, cytidine, or choline nutrients. Data shown are the results from a single experiment with three replicates per condition (three different wells, same 96-well plate). Independent repetition of the experiment (>3, different plates, different days) showed similar results. B) TLC run of C-14 acetate labeled lipids extracted from the different strains. Figure 4C summarizes this experiment.

**Extended Data Figure 2:**
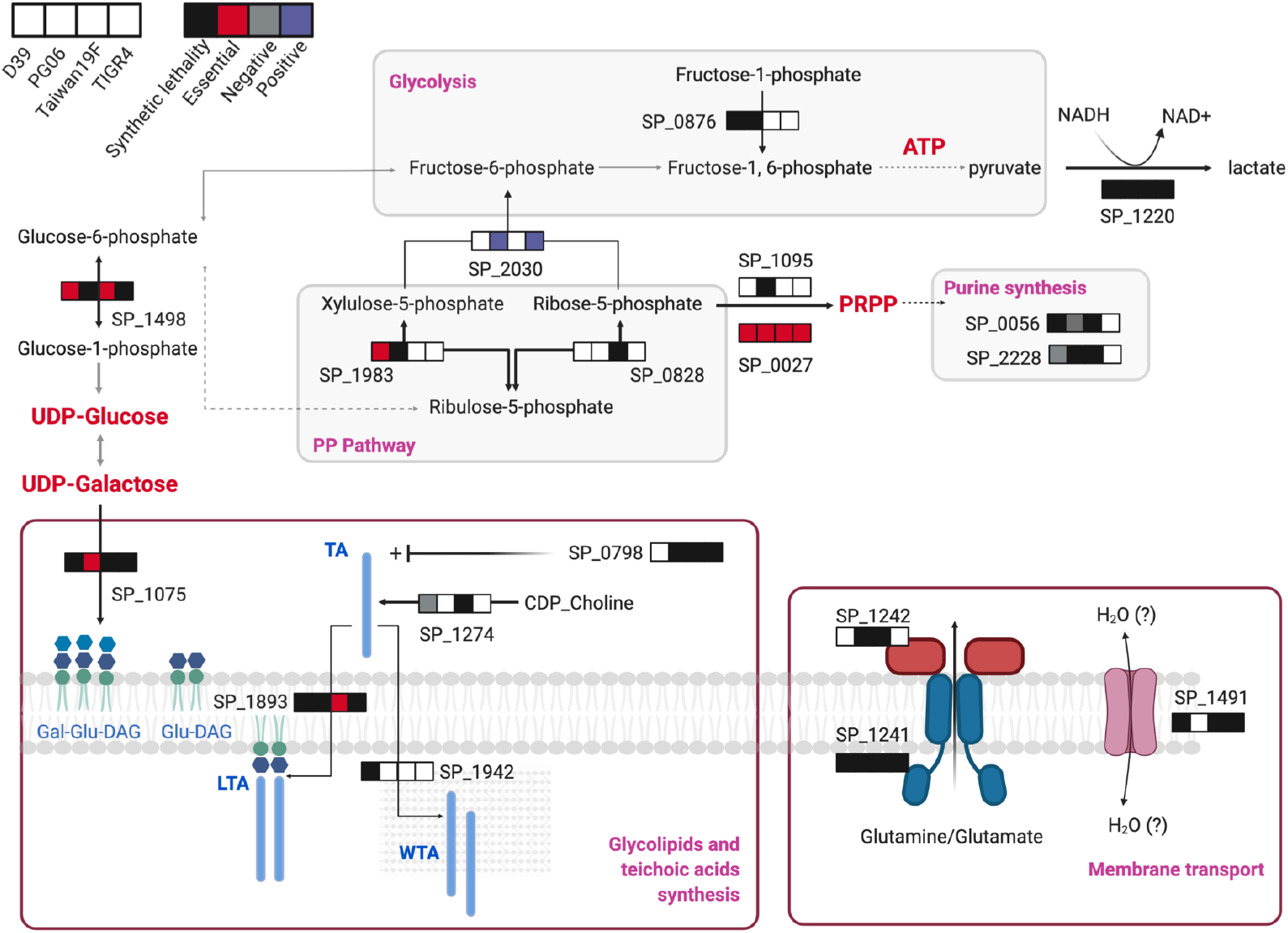
Genes and pathways with genetic interactions with capsule sugar transferase genes. Selection of genetic interactions identified by Tn-Seq using libraries constructed in sugar transferase knockouts derived from four different strains. The order of the squares in the strips indicates the strain and color the type of interaction identified. White square means no interaction. Red bold larger font indicates metabolites that accumulate in the context of *S. pneumoniae* unencapsulated strains ^81^. Gal-Glu-DAG: galactosyl-glucosyl-diacylglycerol, Glu-DAG: glucosyl-diacylglycerol, LTA: lipoteichoic acid, WTA: wall teichoic acid.

**Extended Data Figure 3:**
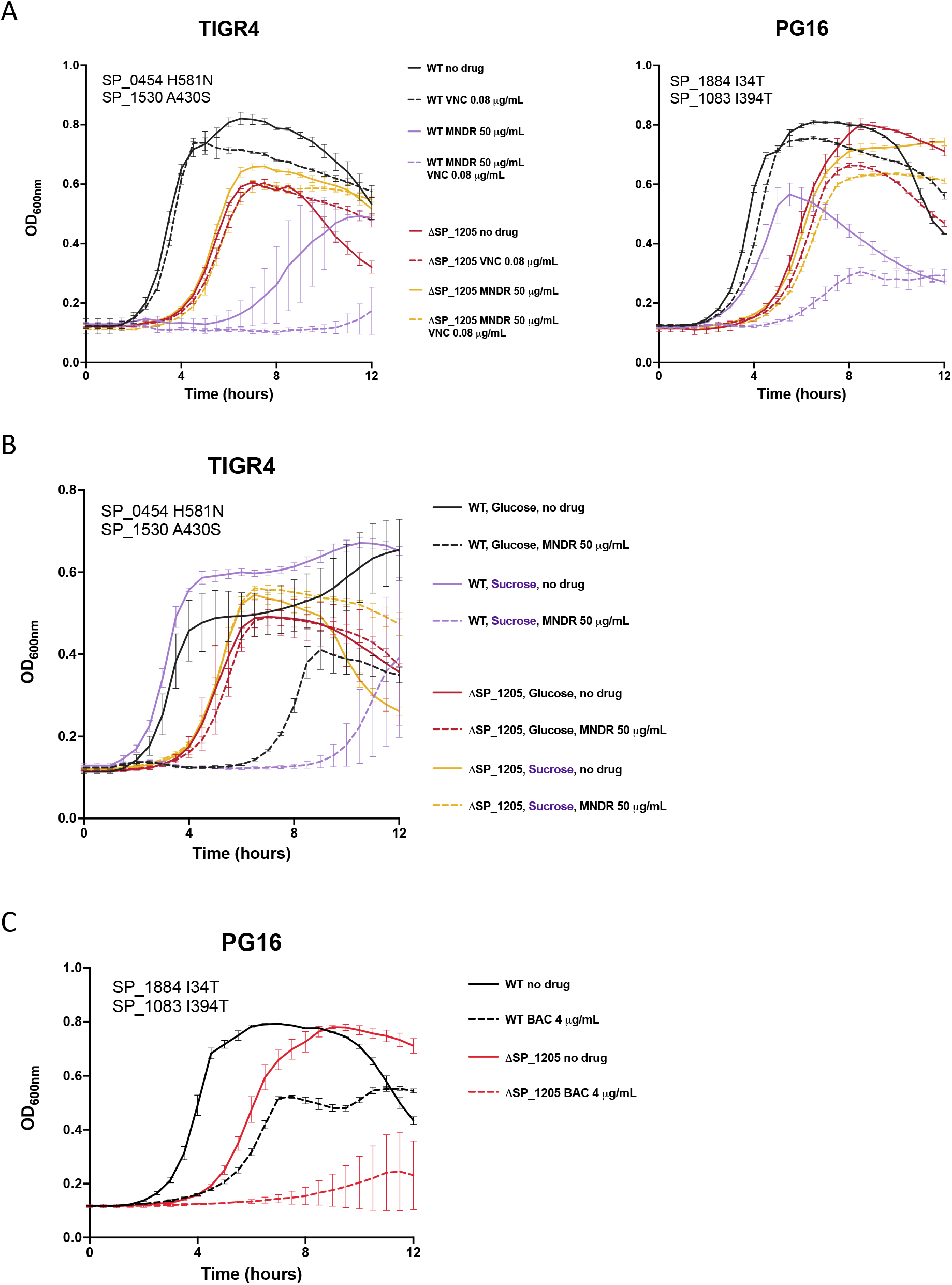
SP_1205 knockouts characterization. A) Synergism between minodronate (MNDR) and vancomycin. Growth phenotype of wild type strains TIGR4 and PG16 (SP_1205 is essential in both) and their derived SP_1205 knockouts in medium SDMM in the presence of non-inhibitory concentrations of vancomycin (VNC), in the presence or absence of the SP_1205 targeting drug minodronate (MNDR), or both. Panels show the suppressive mutations identified in the knockouts. B) Carbon source and SP_1205 requirement. Growth phenotype of wild type strains TIGR4 (SP_1205 is essential) and the derived SP_1205 knockout in medium SDMM with glucose or sucrose as carbon source, and with or without MNDR. Panel shows the suppressive mutations identified in the knockout. C) SP_1205 knockouts bacitracin sensitivity. Growth phenotype of wild type strains PG16 (SP_1205 is essential), and the derived SP_1205 knockout in medium SDMM in the presence or absence of the UPP recycling inhibitory drug bacitracin (Bac). Data from A, B, and C are the results from single experiments with three replicates per condition (three different wells, same 96-well plate). Independent repetition of the experiment (>3) showed similar results.

